# UPF3A and UPF3B are redundant and modular activators of nonsense-mediated mRNA decay in human cells

**DOI:** 10.1101/2021.07.07.451444

**Authors:** Damaris Wallmeroth, Volker Boehm, Jan-Wilm Lackmann, Janine Altmüller, Christoph Dieterich, Niels H. Gehring

**Affiliations:** Institute for Genetics, University of Cologne, 50674 Cologne, Germany; Center for Molecular Medicine Cologne (CMMC), University of Cologne, 50937 Cologne, Germany; CECAD Research Center, University of Cologne, Joseph-Stelzmann-Str. 26, 50931 Cologne, Germany; Cologne Center for Genomics (CCG), University of Cologne, 50931 Cologne, Germany; Berlin Institute of Health at Charité – Universitätsmedizin Berlin, Core Facility Genomics, Charitéplatz 1, 10117 Berlin, Germany and Max Delbrück Center for Molecular Medicine in the Helmholtz Association (MDC), Berlin, Germany; Section of Bioinformatics and Systems Cardiology, Department of Internal Medicine III and Klaus Tschira Institute for Integrative Computational Cardiology, Heidelberg University Hospital, 69120 Heidelberg, Germany; DZHK (German Centre for Cardiovascular Research), Partner site Heidelberg/Mannheim, 69120 Heidelberg, Germany

**Keywords:** Nonsense-mediated mRNA decay, gene paralogs, mRNA turnover, UPF3

## Abstract

The paralogous human proteins UPF3A and UPF3B are involved in recognizing mRNAs targeted by nonsense-mediated mRNA decay (NMD). While UPF3B has been demonstrated to support NMD, contradicting reports describe UPF3A either as an NMD activator or inhibitor. Here, we present a comprehensive functional analysis of UPF3A and UPF3B in human cells using combinatory experimental approaches. Overexpression or knockout of UPF3A as well as knockout of UPF3B did not detectably change global NMD activity. In contrast, the co-depletion of UPF3A and UPF3B resulted in a marked NMD inhibition and a transcriptome-wide upregulation of NMD substrates, demonstrating a functional redundancy between both NMD factors. Although current models assume that UPF3 bridges NMD-activating exon-junction complexes (EJC) to the NMD factor UPF2, UPF3B exhibited only slightly impaired NMD activity in rescue experiments when UPF2 or EJC binding was impaired. Further rescue experiments revealed partially redundant functions of UPF3B domains in supporting NMD, involving both UPF2 and EJC interaction sites and the central region of UPF3. Collectively, UPF3A and UPF3B serve as fault-tolerant NMD activators in human cells.

## Introduction

Precisely regulated expression of correct gene products is indispensable for eukaryotic life. This is underlined by the existence of several quality control mechanisms for gene expression, one of which is the nonsense-mediated mRNA decay (NMD). NMD is primarily known for its ability to eliminate mature mRNAs that contain a premature termination codon (PTC). Thereby, NMD prevents the synthesis and accumulation of C-terminally truncated proteins, which may possess undesirable and potentially disease-causing properties (Frischmeyer & Dietz, 1999). Although the removal of PTC-containing mRNAs was initially considered the most important function of NMD, later studies showed that NMD plays an important role in the post-transcriptional regulation of a substantial part of the transcriptome (He *et al*., 2003; Lelivelt & Culbertson, 1999; Mendell *et al*., 2004; Rehwinkel *et al*., 2005). The importance of the factors involved in NMD is underscored by the severe impact that mutations in components of this machinery have on development in metazoans, up to causing embryonic lethality in mammals (Hwang & Maquat, 2011; Li *et al*., 2015; McIlwain *et al*., 2010; Medghalchi *et al*., 2001; Metzstein & Krasnow, 2006; Weischenfeldt *et al*., 2008; Wittkopp *et al*., 2009).

The final step of gene expression is the cytoplasmic translation of the mRNA by ribosomes. Previous studies suggested that prolonged ribosome stalling at a termination codon indicates improper translation termination and thereby triggers NMD (Amrani *et al*., 2004; Peixeiro *et al*., 2012). This could be caused by a long 3’ untranslated region (UTR) that increases the distance between the stalled ribosome and the poly(A)-binding protein (PABPC1), which normally promotes proper translation termination (Amrani *et al*., 2004). Alternatively, NMD can also be activated by any PTC located more than 50-55 nt upstream of the 3’-most exon-exon junction. Transcripts with such a PTC may be transcribed from mutant genes with nonsense mutations but could also be generated by defective or alternative splicing (Kervestin & Jacobson, 2012). The aforementioned 50-55 nt boundary between NMD-activating and NMD-resistant PTCs is determined by the RNA-binding exon junction complex (EJC), which is deposited by the spliceosome 20-24 nt upstream of every spliced exon-exon junction (Le Hir *et al*., 2000). The EJCs remain attached on the mature mRNA during export into the cytoplasm, where they are removed by translating ribosomes or the disassembly factor PYM1 (Dostie & Dreyfuss, 2002; Le Hir *et al*., 2000). If translation terminates prematurely due to the presence of a PTC, EJCs bound downstream of the PTC serve as a marker for the NMD machinery and the initial activation of NMD (Kim *et al*., 2001; Le Hir *et al*., 2001).

Extensive research over many decades has resulted in a model for EJC-dependent NMD. According to this model, the central factor UPF1 is bound non-specifically to all present transcripts in the cell and is removed from the coding sequence by translating ribosomes (Hogg & Goff, 2010; Hurt *et al*., 2013; Kurosaki & Maquat, 2013; Zund *et al*., 2013). If translation terminates prematurely, UPF1 interacts with the stalled ribosome and serves as the anchoring point for the other NMD factors. The presence of a downstream EJC is detected by the interaction of UPF1 with UPF2, which in turn binds to UPF3. The latter can bind directly to the EJC, resulting in a bridged connection of UPF1 to the EJC (Chamieh *et al*., 2008; Kim *et al*., 2001; Le Hir *et al*., 2001; Weng *et al*., 1996). This series of interactions marks the termination codon as premature and stimulates the phosphorylation of N- and C-terminal SQ motif-containing regions of UPF1 by the kinase SMG1 (Yamashita *et al*., 2001). In its phosphorylated state UPF1 recruits the heterodimer SMG5-SMG7 and/or SMG6, which are responsible for both exoribonucleolytic and endoribonucleolytic degradation of the mRNA, respectively (Boehm *et al*., 2021; Chen & Shyu, 2003; Lejeune *et al*., 2003). The endonuclease SMG6 cleaves the mRNA in close proximity to the PTC, resulting in two mRNA fragments (Eberle *et al*., 2009) of which the 3’ fragment is degraded by the 5’-to-3’ exoribonuclease XRN1 (Eberle *et al*., 2009; Huntzinger *et al*., 2008).

The protein UPF3 plays an important role in the NMD pathway. As detailed above, its main function is believed to bridge the NMD machinery to the EJC providing a physical link between UPF2 and the EJC (Chamieh *et al*., 2008; Kashima *et al*., 2006; Lykke-Andersen *et al*., 2000; Serin *et al*., 2001). With its conserved RNA recognition motif (RRM) in the N-terminus, UPF3 can interact with the C-terminal MIF4G (middle portion of eIF4G) domain of UPF2 (Kadlec *et al*., 2004). The association with the EJC-binding site, formed by the core components EIF4A3, MAGOHB and RBM8A, is enabled by a C-terminal sequence referred to as EJC binding motif (EBM) (Buchwald *et al*., 2010; Gehring *et al*., 2003; Kim *et al*., 2001).

In vertebrates two genes encode for two UPF3 paralogues: UPF3A and UPF3B, each of which expressing two different isoforms generated by alternative splicing (Lykke-Andersen *et al*., 2000; Serin *et al*., 2001). Both human UPF3 proteins contain the same important domains, but differ in details regarding their interactions. According to previous studies, UPF3A and UPF3B are in constant competition for their binding partner UPF2. However, UPF3B binds tighter to UPF2 than UPF3A and therefore UPF3A gets destabilized, and its protein levels are barely detectable under normal conditions (Chan *et al*., 2009).

Recently, UPF3B was reported to interact with the eukaryotic release factor 3 (eRF3) via the so far uncharacterized middle domain (amino acids (aa) 147-256) (Neu-Yilik *et al*., 2017). Due to this interaction and binding of the terminating ribosome, it can delay translation termination, which is known to define aberrant termination events and trigger NMD (Amrani *et al*., 2004; Neu-Yilik *et al*., 2017; Peixeiro *et al*., 2012).

Due to the different molecular characteristics of the UPF3 paralogs, their exact role in NMD is a long-discussed topic. On the one hand, previous studies showed that UPF3A and UPF3B both trigger degradation of a reporter construct when tethered downstream of a termination codon. Notably, the efficiency of UPF3A to elicit NMD was weaker in comparison to UPF3B, which was attributed to a weaker interaction with the EJC (Kunz *et al*., 2006; Lykke-Andersen *et al*., 2000). This would suggest that, at least with respect to their NMD activity, UPF3A and UPF3B serve a similar, perhaps even redundant, function. This notion is supported by the observation that in patients with mutated UPF3B the amount of stabilized UPF3A inversely correlated with the severity of the patients’ clinical phenotypes (Nguyen *et al*., 2012). On the other hand, it was recently reported that loss of UPF3A results in increased transcript destabilization, and UFP3A overexpression in NMD inhibition (Shum *et al*., 2016). This would rather indicate opposing functions of the two UPF3 paralogs with UPF3A being an antagonist of UPF3B and broadly acting as an NMD inhibitor.

In this study, we resolved the controversy about the functions of UPF3A and UPF3B in the NMD pathway using different UPF3 overexpression and knockout (KO) HEK293 cell lines. We found that neither overexpression nor genomic KO of UPF3A resulted in substantial changes of NMD activity or global alterations of the transcriptome. In UPF3B KO cells UPF3A protein levels were upregulated, but NMD activity was maintained at almost normal level. In contrast, the co-depletion of both UPF3 paralogs resulted in a marked NMD inhibition and a global upregulation of PTC-containing transcripts. Moreover, rescue experiments revealed that UPF3 proteins have additional functions besides bridging the EJC and the NMD machinery. Taken together, our data support a model of human NMD, in which UPF3A and UPF3B can replace each other and therefore perform redundant functions.

## RESULTS

### UPF3A overexpression or knockout does not affect NMD efficiency

Prior work using different mammalian models and various experimental approaches reached different conclusions regarding the function of UPF3A in NMD (Fig 1A). Therefore, we set out to re-examine the role of UPF3A in human cells by specifically manipulating its expression levels. Under regular conditions UPF3A is barely present in cultured cells, presumably due to its lower binding affinity to the stabilizing interaction partner UPF2 compared to UPF3B, resulting in a rapid turnover of “free” UPF3A (Chan *et al*., 2009). We hypothesized that increasing the abundance of UPF3A should lead to the stabilization of NMD targets if UPF3A is an NMD inhibitor. To test this hypothesis, we generated Flp-In T-REx 293 (HEK293) cells inducibly overexpressing FLAG-tagged wildtype UPF3A to high protein levels (Fig 1B). Global analysis of the transcriptome using RNA-seq (Fig EV1A and Datasets EV1-EV3) revealed, except for UPF3A itself, no significant differential gene expression (DGE), differential transcript usage (DTU) or alternative splicing (AS) events upon UPF3A overexpression compared to control conditions (Figs 1C and D). Using these RNA-seq data, we analyzed NMD targets that were previously described to be strongly upregulated in UPF3A overexpressing HeLa cells (Shum *et al*., 2016). The DGE analysis and visualization of the respective read coverage showed no substantial effects in our setup (Figs 1E and EV1B-E). Furthermore, quantification of differential transcript usage via IsoformSwitchAnalyzeR (Vitting-Seerup & Sandelin, 2019) could neither detect any differences in the global isoform fraction distribution, nor an accumulation of PTC-containing transcripts (Fig EV1F). Collectively, these analyses indicated that UPF3A overexpression in HEK293 cells does not negatively affect gene expression in general or NMD in particular.

**Figure 1.**
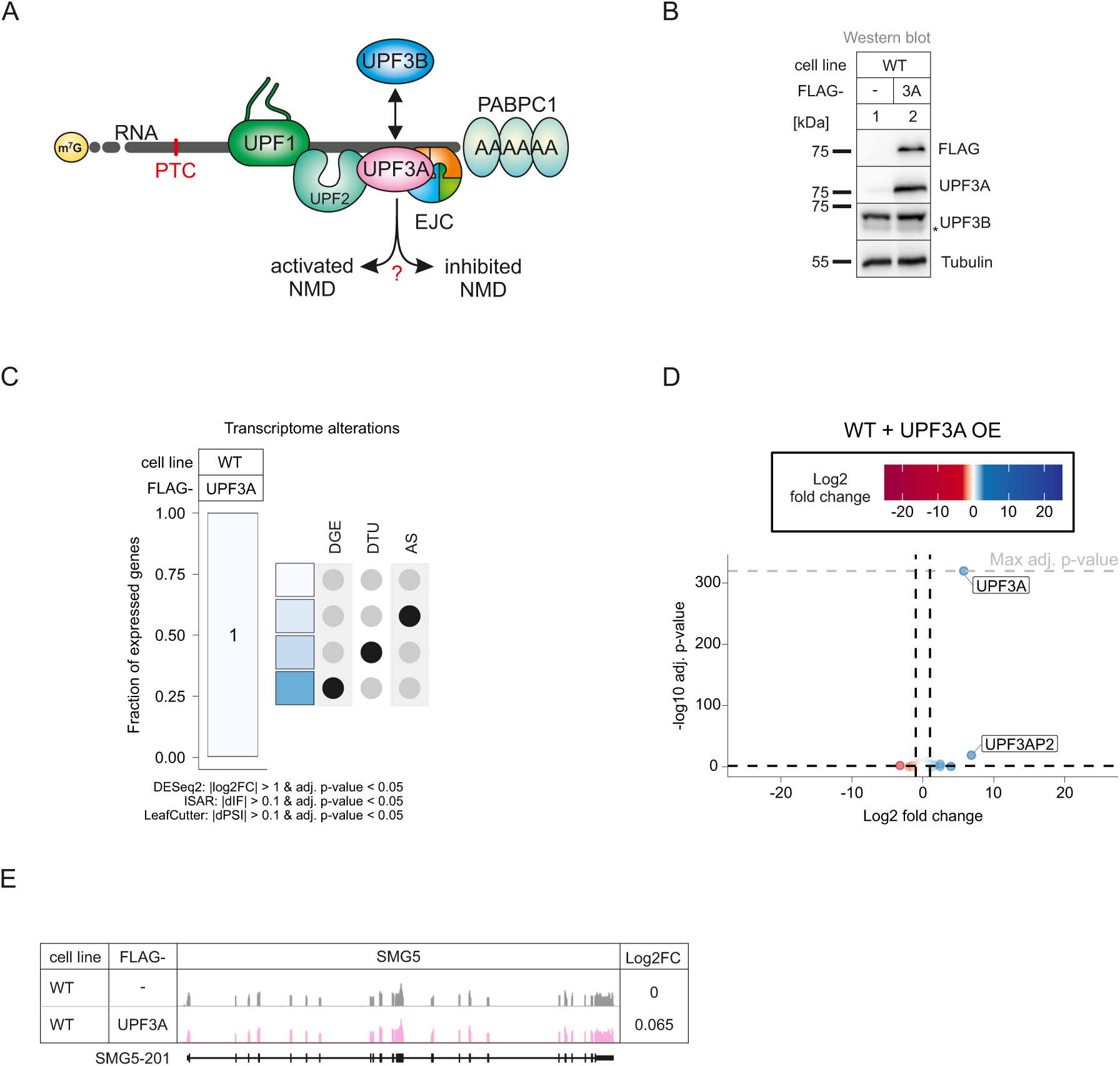
UPF3A overexpression does not affect NMD. **A** Schematic representation of the bridge between UPF1 and the EJC during NMD. Binding of UPF3Ainstead of the stronger bound UPF3B is discussed to either activate or inhibit NMD. **B** Western blot analyses after induced expression of FLAG-tagged UPF3Ain WT HEK 293 cells. Tubulin serves as control. The asterisk indicates unspecific bands. **C** Fraction of expressed genes (genes with non-zero counts in DESeq2) were calculated which exhibit individual or combinations of differential gene expression (DGE), differential transcript usage (DTU) and/or alternative splicing (AS) events in WT cells overexpressing UPF3A using the respective computational analysis (cutoffs are indicated). AS and DTU events were collapsed on the gene level. For DGE, p-values were calculated by DESeq2 using a two-sided Wald test and corrected for multiple testing using the Benjamini-Hochberg method. For DTU, p-values were calculated by IsoformSwitchAnalyzeR (ISAR) using a DEXSeq-based test and corrected for multiple testing using the Benjamini-Hochberg method. For AS, p-values were calculated by LeafCutter using an asymptotic Chi-squared distribution and corrected for multiple testing using the Benjamini-Hochberg method. **D** Volcano plot showing the differential gene expression analyses from the RNA-Seq dataset of WT cells overexpressing UPF3A. The log2 fold change is plotted against the -log10 adjusted p-value (adj. p-value). P-values were calculated by DESeq2 using a two-sided Wald test and corrected for multiple testing using the Benjamini-Hochberg method. OE = overexpression. **E** Read coverage of SMG5 from WT HEK 293 RNA-seq data with or without induced UPF3A overexpression shown as Integrative Genomics Viewer (IGV) snapshot. Differential gene expression (from DESeq2) is indicated as Log2 fold change (Log2FC) on the right. Schematic representation of the protein coding transcript below.

Next, we approached the question of UPF3A function in the opposite way by generating UPF3A knockout (KO) HEK293 cell lines. Using CRISPR-Cas9 genome editing we isolated three clones that lacked the UPF3A-specific band on the Western blot even after downregulation of UPF3B (Fig 2A). Two clones (14 and 20) were characterized in detail.

**Figure 2.**
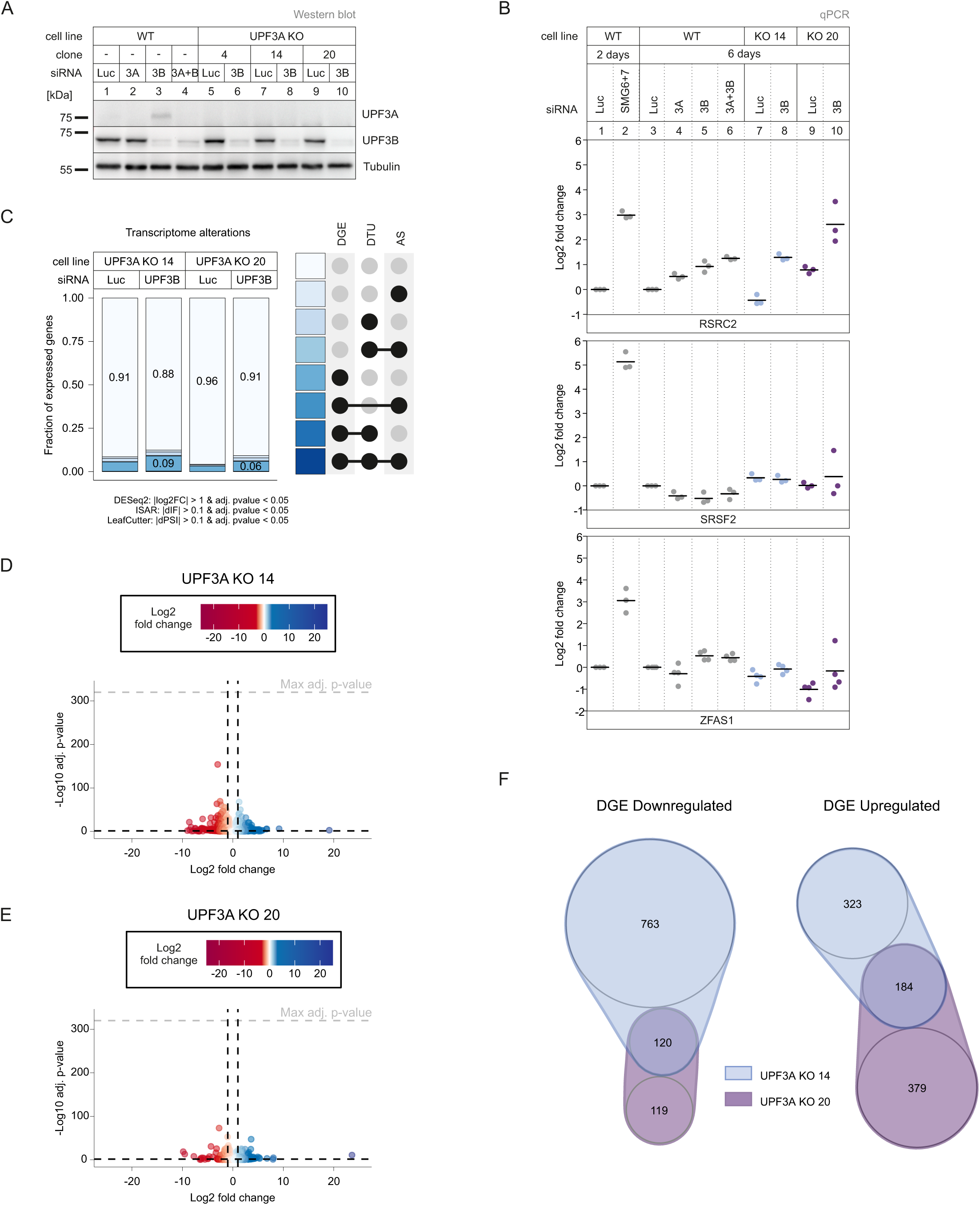
UPF3A KOs show light NMD-independent transcriptome alterations. **A** Western blot analysis of WT and UPF3AKO cells (clones 4, 14 and 20) with the indicated siRNAtreatments. UPF3Aand UPF3B protein levels were detected, Tubulin serves as control. **B** Quantitative RT-PCR of the indicated cell lines treated with the indicated siRNAs for 2 or 6 days. For RSRC2 and SRSF2 the ratio of NMD isoform to canonical isoform was calculated. ZFAS1 expression was normalized to C1orf43 reference. Data points and means are plotted as Log2 fold change (n=3 for RSRC2 and SRSF2, n=4 for ZFAS1). **C** Fraction of expressed genes (genes with non-zero counts in DESeq2) were calculated which exhibit individual or combinations of differential gene expression (DGE), differential transcript usage (DTU) and/or alternative splicing (AS) events in the indicated conditions using the respective computational analysis (cutoffs are indicated). AS and DTU events were collapsed on the gene level. For DGE, p-values were calculated by DESeq2 using a two-sided Wald test and corrected for multiple testing using the Benjamini-Hochberg method. For DTU, p-values were calculated by IsoformSwitchAnalyzeR using a DEXSeq-based test and corrected for multiple testing using the Benjamini-Hochberg method. For AS, p-values were calculated by LeafCutter using an asymptotic Chi-squared distribution and corrected for multiple testing using the Benjamini-Hochberg method. **D,E** Volcano plots showing the differential gene expression analyses from the indicated RNA-Seq datasets (D UPF3A KO clone 14, E UPF3A KO clone 20). The log2 fold change is plotted against the -log10 adjusted p-value (padj). P-values were calculated by DESeq2 using a two-sided Wald test and corrected for multiple testing using the Benjamini-Hochberg method **F** nVenn Diagram showing the overlap of up- or downregulated genes in the UPF3A KO cell lines 14 and 20. Log2 fold change <1 (downregulated) or >1 (upregulated) and adjusted p-value (padj) < 0.05. DGE = Differential Gene Expression.

In both cell lines, the UPF3A genomic locus contained insertions and/or deletions causing frame-shifts and eventually PTCs (Figs EV2A-C). To gain a first impression of the NMD activity in the UPF3A KO cells, the transcript levels of three known exemplary endogenous NMD targets, RSRC2, SRSF2 and ZFAS1 were determined by qPCR (Boehm *et al*., 2021; Lykke-Andersen *et al*., 2014; Sureau *et al*., 2001). WT HEK293 cells treated with SMG6 and SMG7 siRNAs were used as a positive control for severe NMD inhibition (Fig 2B)(Boehm *et al*., 2021). While the absence of UPF3A did not result in abundance changes of SRSF2 and RSRC2 NMD-sensitive isoforms (mean log2FC between -0.44 and 0.79), ZFAS1 mRNA levels were slightly decreased in the UPF3A KO cells compared to WT cells (mean log2FC -0.42 and -1.01 for UPF3A KO clones 14 and 20, respectively; Fig 2B). However, this effect was not rescued by the (over)expression of transgenic UPF3A, indicating that it is not caused by the lack of UPF3A but rather represents random variations in gene expression or clonal effects (Figs EV2D-E). To get a complete overview of the effects of the UPF3A KO, we performed RNA-seq of two UPF3A KO cell lines with or without an additional UPF3B knockdown (KD; Fig EV2F and Datasets EV1-EV3). Initially, we focused on the UPF3A KO cell lines without KDs, for which the global transcriptome analysis revealed that about 4-9 % of the expressed genes are altered (Figs 2C-E). The observation that in the absence of UPF3A more genes were downregulated than upregulated (1002 vs. 886) could be an indicator for UPF3A NMD inhibiting properties (Fig 2F). However, the majority of genes with altered expression were clone specific and only 120 genes showed downregulation in both UPF3A KO cell lines (Fig 2F). Investigation of selected targets that were significantly up- or downregulated in both clones revealed that the changes were not rescued after UPF3A overexpression, suggesting that they are UPF3A-independent (Figs EV2D-E). Another indication that UPF3A depletion does not generally affect NMD efficiency came from the DTU analysis.

Although clone 14 showed a minor downregulation of PTC-containing transcripts, which could indicate more active NMD, this effect was not reproducible in the second clone (Fig EV2G). In view of and in combination with the results shown in Fig 1, this strongly suggests that neither the overexpression nor the depletion of UPF3A substantially alters (negatively or positively) the efficiency of NMD. In conclusion, our data argue against a role for UPF3A as a general negative NMD regulator in human cell lines.

### NMD is functional in the absence of UPF3B

Next, we investigated the RNA-seq data of UPF3B knockdowns in the UPF3A KO cells (Datasets EV1-EV3). We observed that this combination resulted in more transcriptome alterations and an increase of PTC-containing isoforms (Figs 2B-C, EV2H and I). Although the UPF3B KD alone could be in principle responsible for this effect, the results could also be an indicator for redundant functions of the two UPF3 paralogs. To explore this hypothesis, we decided to generate UPF3B KO cell lines. Western blot analysis demonstrated that a UPF3B KD is less efficient in reducing the produced protein than the UPF3B KO in the two clones designated as 90 and 91 (Figs 3A and EV3A-B). In addition, we observed a strong upregulation of UPF3A after depletion (KO) or reduction (KD) of UPF3B, as described before (Chan *et al*., 2009). In the absence of UPF3B no changes in the expression of the respective NMD-sensitive isoforms of the NMD-targets RSRC2 and SRSF2 was observed (Fig 3B). This indicates that either UPF3B is not essential for NMD or that another protein is able to compensate for its loss. The most obvious candidate for this function is its own paralog UPF3A, which was also suggested previously to functionally replace UPF3B in NMD. Indeed, knocking down UPF3A in the UPF3B KO cells resulted in the increase of NMD-sensitive RSRC2 and SRSF2 isoforms (Fig 3B). Of note, the combination of the UPF3B KO with UPF3A KD showed stronger effects than the previously analyzed UPF3A KO plus UPF3B KD. This is probably caused by the lower KD efficiency of the UPF3B siRNAs which can be observed by comparing the respective protein levels (Fig 2A vs. Fig 3A). We suspect that the remaining UPF3B levels after siRNA-mediated UPF3B KD still support NMD.

**Figure 3.**
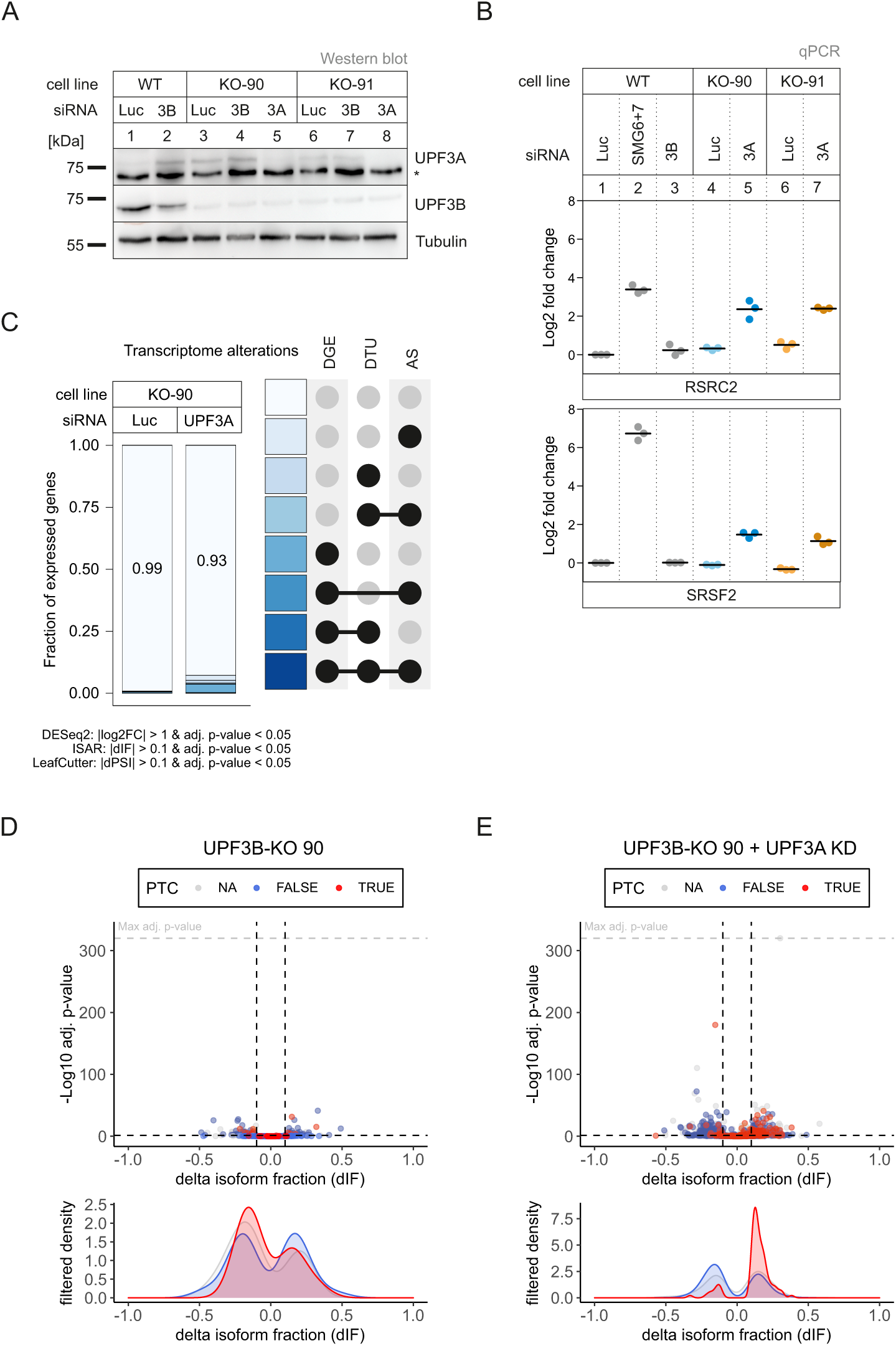
Loss of UPF3B does not affect NMD efficiency, only in combination with KD of UPF3A. **A** Western blot analysis of WT and UPF3B KO cells (clones 90 and 91) combined with the indicated knockdowns. UPF3A and UPF3B (AK-141) protein levels were detected, Tubulin serves as control. The asterisk indicates unspecific bands. **B** Quantitative RT-PCR of the indicated cell lines with the indicated knockdowns. For RSRC2 and SRSF2 the ratio of NMD isoform to canonical isoform was calculated. Data points and means are plotted as Log2 fold change (n=3). **C** Fraction of expressed genes (genes with non-zero counts in DESeq2) were calculated which exhibit individual or combinations of differential gene expression (DGE), differential transcript usage (DTU) and/or alternative splicing (AS) events in the indicated conditions using the respective computational analysis (cutoffs are indicated). AS and DTU events were collapsed on the gene level. For DGE, p-values were calculated by DESeq2 using a two-sided Wald test and corrected for multiple testing using the Benjamini-Hochberg method. For DTU, p-values were calculated by IsoformSwitchAnalyzeR using a DEXSeq-based test and corrected for multiple testing using the Benjamini-Hochberg method. For AS, p-values were calculated by LeafCutter using an asymptotic Chi-squared distribution and corrected for multiple testing using the Benjamini-Hochberg method. **D,E** Volcano plots showing the differential transcript usage (via IsoformSwitchAnalyzeR) in various RNA-Seq data. Isoforms containing GENCODE (release 33) annotated PTC (red, TRUE), regular stop codons (blue, FALSE) or having no annotated open reading frame (grey, NA) are indicated. The change in isoform fraction (dIF) is plotted against the -log10 adjusted p-value (adj.p-value). Density plots show the distribution of filtered isoforms in respect to the dIF, cutoffs were |dIF| > 0.1 and adj. p-value < 0.05. P-values were calculated by IsoformSwitchAnalyzeR using a DEXSeq-based test and corrected for multiple testing using the Benjamini-Hochberg method.

To gain more transcriptome-wide information, we performed RNA-seq of the UPF3B KO clone 90, with and without UPF3A siRNA treatment (Fig EV3C and Datasets EV1-EV3). The global effects detected in the RNA-seq data correlated well with the NMD inhibition seen for single targets (Figs 3C and EV3D-G). Analysis of the differential transcript usage revealed an upregulation of transcripts annotated with a PTC only in the UPF3B KO cells with additionally downregulated UPF3A (Figs 3D and E). This increase of NMD-sensitive transcripts could not be observed in the plain UPF3B KO cells with UPF3A naturally upregulated. All together this data strongly suggested at least partial redundancy of UPF3A and UPF3B, since KO of only one paralog was irrelevant for NMD functionality. Furthermore, we were able to show that also in the absence of UPF3B an overexpression of UPF3A has no negative effects on NMD efficiency, supporting the previous conclusion of UPF3A not being a negative NMD regulator (Figs EV3H and I).

### Stronger NMD impairment in UPF3A-UPF3B double KO cells

Considering that UPF3A and UPF3B seemingly carry out redundant functions, we decided to create a cell line completely lacking both paralogs and aimed to generate UPF3A-UPF3B double KO cells (UPF3 dKO). These cells should show stronger effects than the combination of a KO and a KD, since residual amounts of protein were typically still detected after siRNA treatment. Using the UPF3B-KO clone 90 as parental cell line, two potential UPF3 dKO clones 1 and 2 were generated (Fig 4A), which differed in the guide RNAs used to target exon 1 of UPF3A. We confirmed that both cell lines contained frame shift-inducing insertions/deletions at the respective positions in the UPF3A gene (Figs 4B, EV4A and B). We first explored by qPCR how strongly the dKO affected NMD (Fig 4C). For all three tested genes, the expression of the NMD-sensitive isoform was further increased compared to the previously used combination of UPF3B KO with additional UPF3A KD. Of note, the NMD inhibitory effect observed in the dKOs became more pronounced after UPF3B siRNA transfection, suggesting that low levels of residual UPF3B protein were still present in the dKO cells (Fig EV4C).

**Figure 4.**
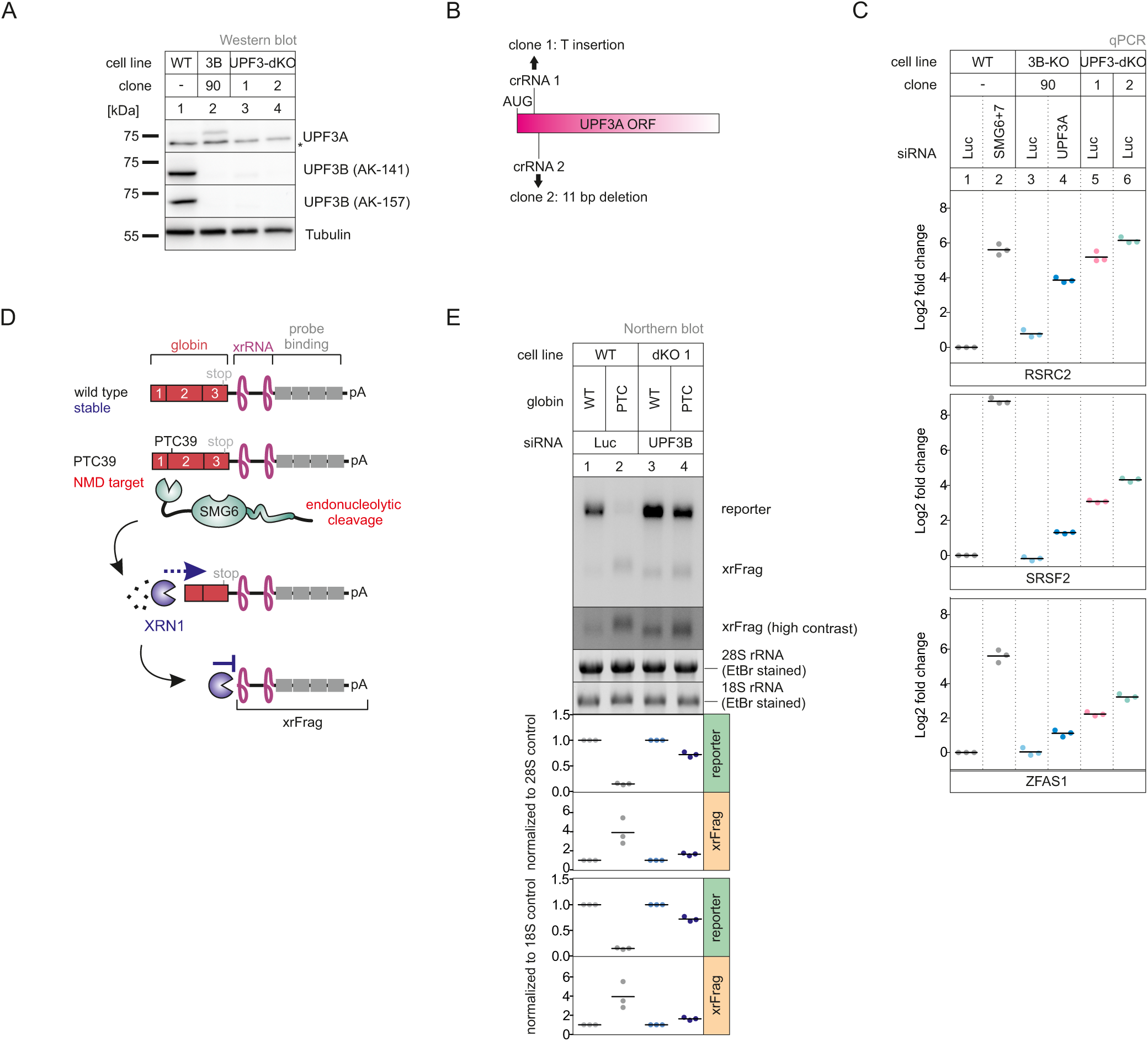
KO of both UPF3 paralogs results in strongly impaired NMD. **A** Western blot analysis of WT, UPF3B KO and UPF3A-UPF3B doubleKO cells (clones 1 and 2). UPF3Aand UPF3B protein levels were detected, Tubulin serves as control. The asterisk indicates unspecific bands. **B** Schematic depiction of the insertion/deletion in the UPF3Aopen reading frame resulting in the additional UPF3AKO in the UPF3B KO clone 90 generating UPF3 dKO clones. **C** Quantitative RT-PCR of the indicated samples with the indicated KDs. For RSRC2 and SRSF2 the ratio of NMD isoform to canonical isoform was calculated. ZFAS1 expression was normalized to C1orf43 reference. Data points and means are plotted as Log2 fold change (n=3). **D** Schematic overview of the globin reporter constructs and their functional elements. **E** Northern blot analysis of globin reporter and xrFrag. Ethidium bromide stained 28S and 18S rRNAs are shown as controls. Quantification results are shown as data points and mean (n=3).

The expression levels of endogenous NMD substrates could be influenced by transcription rates or other indirect effects, which could lead to over- or underestimating NMD inhibition. Therefore, we investigated the NMD efficiency in the dKO cells using the well-established β-globin NMD reporter. To this end, we stably integrated β-globin WT or PTC39 constructs in WT and UPF3 dKO cells (Fig 4D). These reporters also contained XRN1-resistant sequences (xrRNAs) in their 3’ UTRs, which allowed us to analyze not only the degradation of the full-length reporter mRNA but also to quantify decay intermediates (called xrFrag) (Boehm *et al*., 2016; Voigt *et al*., 2019). The PTC39 mRNA was efficiently degraded and a strong xrFrag observed in WT cells, whereas in dKO cells the PTC39 reporter accumulated to high levels (72% compared to the WT mRNA), which was accompanied by a decrease in the amount of xrFrag. (Fig 4E, lane 2 vs. lane 4). In line with the previous observations, this result indicated a strong decrease of NMD activity upon the KO of both UPF3 paralogs using a robust NMD reporter pair expressed independently of endogenous NMD substrates, supporting the functional redundancy of UPF3A and UPF3B in NMD.

To establish transcriptome-wide insights into UPF3A and UPF3B function, we carried out RNA-seq for both dKO clones, which was combined with and without UPF3B KD treatment to eliminate as many of the potentially present remaining UPF3B proteins (Fig EV5A and Datasets EV1-EV3). Differential gene expression analysis showed that more than three times as many genes were upregulated than downregulated in both dKO cells (Figs 5A and EV5B). This is consistent with the redundant role of UPF3B and UPF3A as supporting NMD factors. The considerable overlap between both clones also suggests that we identified high-confidence UPF3 targets. Furthermore, 1072 of these gene were also significantly upregulated in previously generated SMG7 KO plus SMG6 KD data (Fig EV5C, ref data: (Boehm *et al*., 2021)) indicating that these are universal NMD-targets and not specific to a certain branch of the NMD pathway. In addition to DGE, subsets of the expressed genes showed changes in alternative splicing or/and differential transcript usage (Fig 5B). In total, 14-16 % of the global transcriptome showed single or combined changes (DGE, DTU and/or AS) and up to 20% when the cells were treated with an additional UPF3B KD.

**Figure 5.**
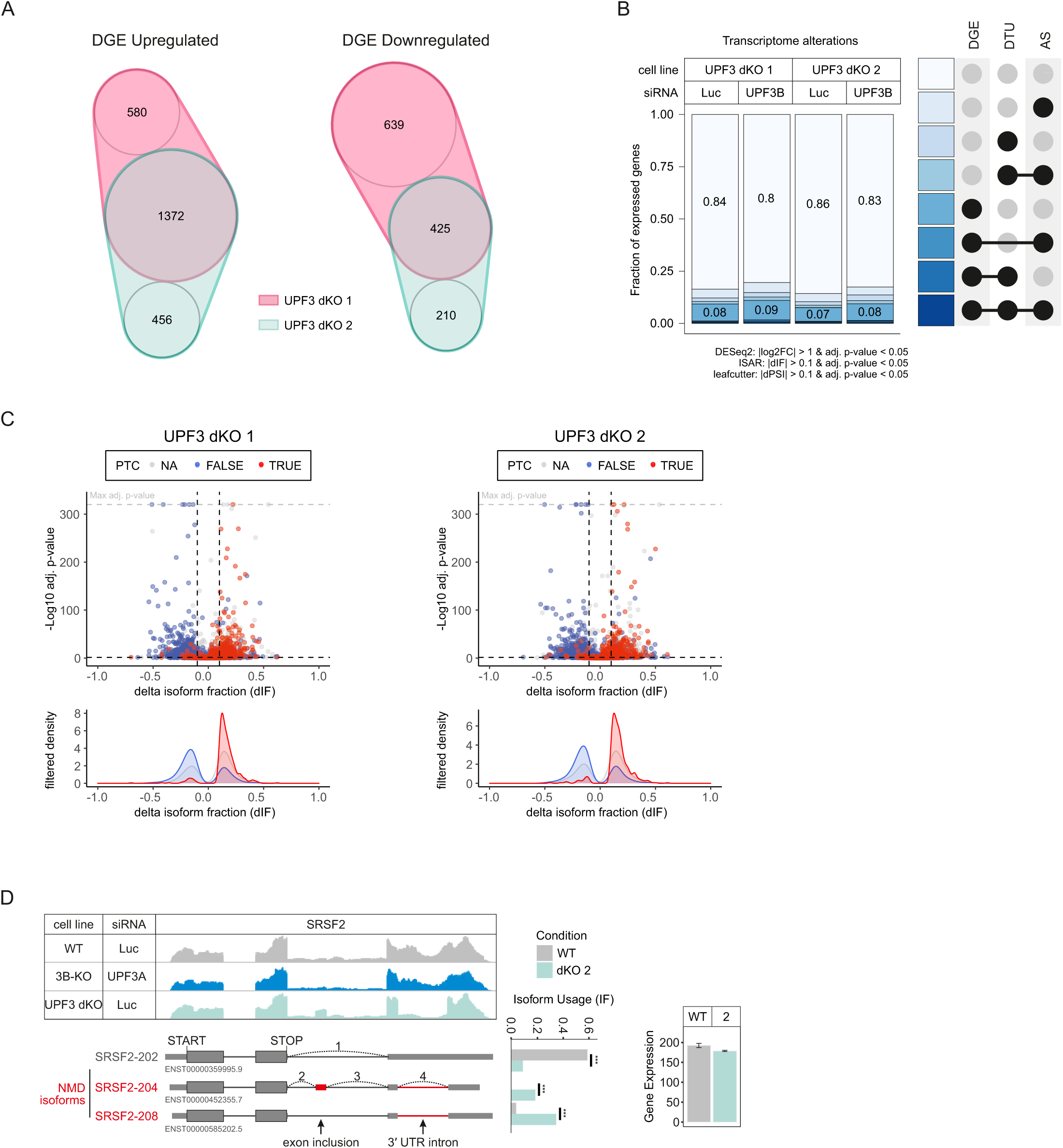
RNA-seq reveals strong global upregulation of NMD-sensitive targets upon UPF3 dKO in HEK293 cells. **A** nVenn Diagram showing the overlap of up- or downregulated genes in the UPF3 dKO cell lines 1 and 2. Log2 fold change <-1 (downregulated) or >1 (upregulated) and adjusted p-value (padj) < 0.05. DGE = Differential Gene Expression. **B** Fraction of expressed genes (genes with non-zero counts in DESeq2) were calculated which exhibit individual or combinations of differential gene expression (DGE), differential transcript usage (DTU) and/or alternative splicing (AS) events in the indicated conditions using the respective computational analysis (cutoffs are indicated). AS and DTU events were collapsed on the gene level. For DGE, p-values were calculated by DESeq2 using a two-sided Wald test and corrected for multiple testing using the Benjamini-Hochberg method. For DTU, p-values were calculated by IsoformSwitchAnalyzeR using a DEXSeq-based test and corrected for multiple testing using the Benjamini-Hochberg method. For AS, p-values were calculated by LeafCutter using an asymptotic Chi-squared distribution and corrected for multiple testing using the Benjamini-Hochberg method. **C** Volcano plots showing the differential transcript usage (via IsoformSwitchAnalyzeR) in various RNA-Seq data. Isoforms containing GENCODE (release 33) annotated PTC (red, TRUE), regular stop codons (blue, FALSE) or having no annotated open reading frame (grey, NA) are indicated. The change in isoform fraction (dIF) is plotted against the -log10 adjusted p-value (adj.p-value). Density plots show the distribution of filtered isoforms in respect to the dIF, cutoffs were |dIF| > 0.1 and adj.p-value < 0.05. P-values were calculated by IsoformSwitchAnalyzeR using a DEXSeq-based test and corrected for multiple testing using the Benjamini-Hochberg method. **D** Read coverage of SRSF2 from the indicated RNA-seq sample data with or without UPF3A siRNA treatment shown as Integrative Genomics Viewer (IGV) snapshot. The canonical and NMD-sensitive isoforms are schematically indicated below. Quantification of isoforms by IsoformSwitchAnalyzeR (right).

In agreement with NMD inhibition in the dKOs, we saw that many transcripts containing a PTC were up-regulated, while the corresponding transcripts without a PTC were down-regulated. (Figs 5C and EV5D). Under these conditions, the IGV snapshot of the NMD-target SRSF2 showed NMD-inducing exon inclusion and 3’ UTR splicing events, which were not visible in combined UPF3B KO/UPF3A KD cells (Fig 5D). Collectively, the RNA-seq data support the previously observed strong NMD inhibition in response to the complete absence of both UPF3 paralogs and hence their proposed redundancy.

### UPF3A supports NMD independent of a bridge function

Next, we aimed to analyze whether the severe effects in the dKOs are at least partly due to the loss of a protein-protein interaction bridge between UPF2 and the EJC, while the presence of UPF3A in the UPF3B KOs preserved this function ensuring NMD functionality. Therefore, we expressed FLAG-tagged UPF2 in WT, UPF3B KO and UPF3A-UPF3B dKO cells and analyzed the UPF2 interactome using mass spectrometry (Dataset EV4). Consistent with the previously described interaction partners, we found many NMD factors as well as EJC proteins in the UPF2 interactome in WT cells (Fig 6A). Contrary to our expectation, the three EJC core components (EIF4A3, RBM8A, MAGOHB) barely co-precipitated with UPF2 in the absence of UPF3B (compared to control: log2 FC = 0.58, 0.63 and 0.74, respectively; Fig 6B) and were therefore strongly decreased in comparison to the WT cells (Fig EV6A). Hence, the UPF2- bound UPF3A was unable to establish a stable interaction with the EJC. Surprisingly, in the UPF3B KO cells the EJC-associated CASC3 still showed relatively high levels of co-precipitation (log2 FC = 4.42), which therefore appears to be independent of the interaction with the other EJC components. In the dKOs all interactions with EJC proteins including CASC3 were completely lost (Figs 6C and EV6B). The latter was also observed in a comparable approach employing stable isotope labeling with amino acids in cell culture (SILAC) to analyze the UPF2 interactome in the WT and dKO cells (Dataset EV5). All EJC core components that were highly co-precipitated in WT cells were lost in the absence of both UPF3 paralogs (Figs EV6C-E).

**Figure 6.**
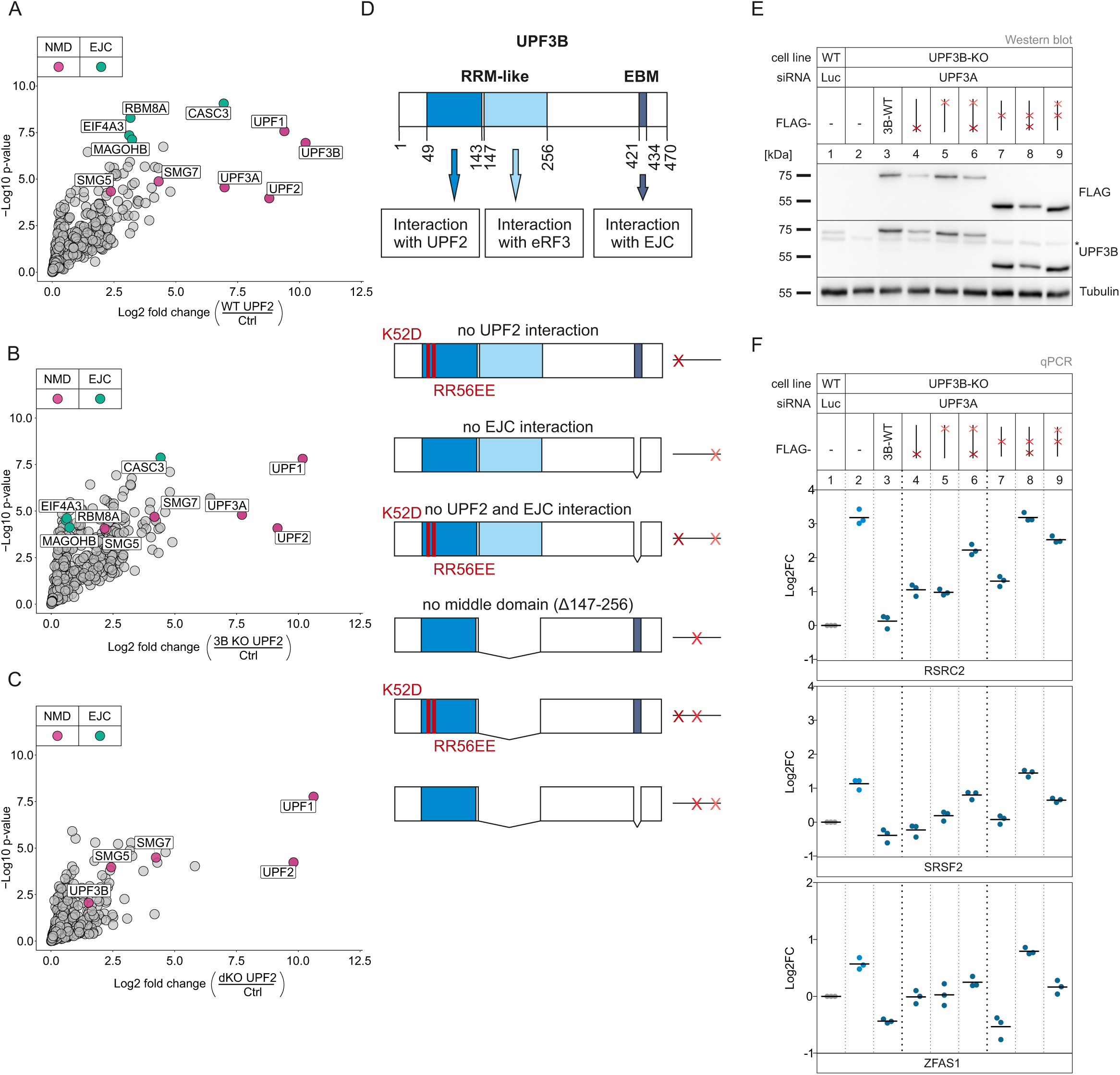
Interaction of UPF3A and UPF3B with the EJC is dispensable to elicit NMD. **A-C** Volcano plots of label free mass spectrometry-based analysis of the interaction partners of UPF2 in WT cells treated with control siRNAs and the UPF3B KO clone 90 and dKO clone 1 both treated with siRNAs targeting UPF3B (n = 4 biologically independent samples). (A) FLAG-UPF2 in WT against FLAG-GST control in WT cells, (B) UPF2 in 3B KO cells against FLAG control in WT cells, (C) UPF2 in dKO cells against FLAG control in WT cells. Points labeled in purple indicate NMD factors; points labeled in turquoise indicate EJC components. Cut offs: Log2 fold change ≥ 0 **D** Schematic representation of the UPF3B protein domains and the respective functions. Below are the mutated rescue constructs and their respective abstract placeholders. **E** Western blot analysis of WT and UPF3B KO clone 90 with Luciferase and UPF3AKDs respectively. Monitored expression of the FLAG-tagged UPF3B rescue construct shown in (D). Rescue construct protein levels were detected with anti-FLAG and anti-UPF3B (AK-141) antibodies. Tubulin serves as control. The asterisk indicates unspecific bands. **F** Quantitative RT-PCR of the samples from (E). For RSRC2 and SRSF2 the ratio of NMD isoform to canonical isoform was calculated. ZFAS1 expression was normalized to C1orf43 reference. Data points and means are plotted as log2 fold change (n=3).

### Partially redundant functions of UPF3B domains are required for NMD

With regard to the surprising observation that UPF3A apparently elicits NMD without interacting with the EJC, we aimed to investigate the molecular features required by UPF3 to support NMD via rescue experiments. In principle, the UPF3 dKO cells represent an ideal system for this approach. However, apart from the residual amounts of UPF3B that were still expressed, we noticed that the UFP3 dKOs were able to upregulate the expression of a shortened UPF3B variant after long-term cultivation.

Since this phenomenon did not occur in the single UPF3B KOs, we generated stable UPF3B KO cell lines expressing various inducible UPF3B constructs with individual or combined binding site mutations (Figs 6D and E and EV6F). Transfection of these cells with UPF3A siRNAs resulted in the robust depletion of UPF3 for the analysis of the rescue activities of individual UPF3B variants. Considering the established role of UPF3B as a bridge between UPF2 and the EJC, which we validated in the mass spec analysis, it was surprising to see that disruption of either of these interactions did not affect or only mildly affected UPF3B’s rescue capacity, depending on the analyzed NMD substrate (Fig 6F, lane 3 vs. lanes 4 and 5). This is partially consistent with the observed functional NMD in UPF3B KOs, despite the apparent inability of UPF3A to form a bridge between UPF2 and the EJC (Figs 3C and 6B). However, mutating both UPF3B binding sites (disrupting UPF2 and EJC binding) largely inactivated the NMD-related function of UPF3B.

It was recently reported that EJC-bound or free UPF3B can interact with the eukaryotic release factor 3 (eRF3) via the so far uncharacterized middle domain (aa 147-256) (Neu-Yilik *et al*., 2017). With this interaction and binding of the terminating ribosome, UPF3B can delay translation termination, which defines aberrant termination events and triggers NMD (Amrani *et al*., 2004; Neu-Yilik *et al*., 2017; Peixeiro *et al*., 2012). To investigate the impact of this interaction on NMD, we created UPF3B variants lacking that specific middle domain or combined the deletion with the previously used interaction mutations (Fig 6D and E). We observed a similar pattern as the UPF3B mutants examined in the previous experiment: when only the middle domain was deleted, UPF3B was able to rescue NMD comparable to the WT protein or slightly worse (Fig 6F lane 7). In combination with a mutation in the UPF2- or the EJC binding site its function in NMD was severely impaired (lanes 8 and 9). This suggests that if the classic bridge formation is inhibited by removing either of the interaction sites, UPF3B relied on the function carried out by the uncharacterized middle domain.

As further support for the NMD promoting effect of UPF3A on NMD, we performed a rescue experiment in the above used UPF3B KO UPF3A KD cells (Figs 7A and B). Expression of the siRNA insensitive UPF3A construct restored NMD functionality with similar efficiency as the UPF3B rescue (Fig 7C lane 3 vs. lane 5), underlining our previous statement: UPF3A supports and elicits NMD comparably to its paralog UPF3B.

**Figure 7.**
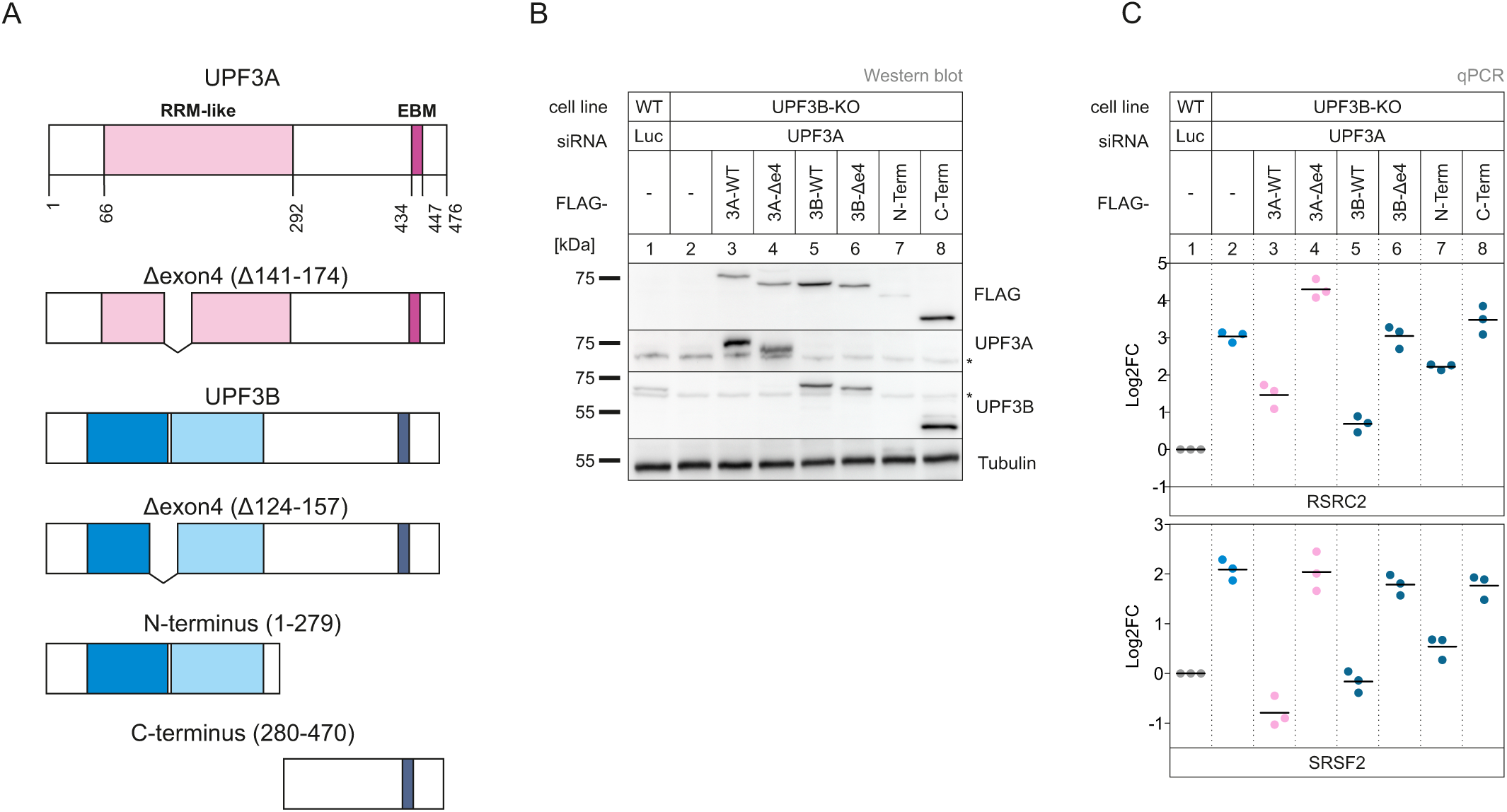
WT UPF3A can rescue NMD in full extent. Deletion on exon 4 disrupts functionality in both paralogs. **A** Schematic representation of the UPF3Aand UPF3B protein domains. Below are the respective mutated rescue constructs. **B** Western blot analysis of WT and UPF3B KO clone 90 with Luciferase and UPF3A KDs respectively. Monitored expression of the FLAG-tagged UPF3A and UPF3B rescue construct shown in (A). Rescue construct protein levels were detected with anti-FLAG, anti-UPF3Aand anti-UPF3B (AK-141) antibodies. Tubulin serves as control. The asterisk indicates unspecific bands. **C** Quantitative RT-PCR of the samples from (B). For RSRC2 and SRSF2 the ratio of NMD isoform to canonical isoform was calculated. Data points and means are plotted as log2 fold change (n=3).

Also comparable to UPF3B, UPF3A has a second naturally occurring isoform but instead of skipping exon 8 (like UPF3B) it excludes its fourth exon. This isoform is transcribed in approximately one third of the cases (Fig EV6G) in WT HEK293 cells but cannot be detected on protein levels. We were interested, whether this isoform was as potent to elicit NMD as the full-length construct. Expression of the exon 4 UPF3A deletion construct in the UPF3 depleted cells showed no rescue (Fig 7C lane 4). Hence, exon 4 must encode for an essential region required for the bridge-independent function of UPF3A.

In view of these observations and the close proximity of exon 4 (124-157) to the middle domain (147-256), we decided to investigate which effect the deletion of the homologous exon 4 in the paralog UPF3B has. The UPF3B Δe4 construct behaved like the corresponding UPF3A construct and showed no NMD rescue activity (lane 6). However, the expression of the N-terminus of UPF3B was able to restore NMD comparably to the WT protein (lane 7). This is consistent with all our previous findings, since the first 279 amino acids contain the UPF2 binding site as well as the middle domain, which was shown to be sufficient to elicit NMD (Fig 6F lane 5). Due to the fact that the C-terminus contains only one interaction site and lacks exon 4, its incapability to rescue NMD is in line with our previous experiments. Overall, our results identify exon 4 of UPF3 as a previously unnoticed region that is essential for its function in NMD.

## Discussion

Methodological advances - be it improved analytics or novel experimental approaches - can help to find new answers to old biological problems. Equipped with powerful new molecular biology methods, we set out to answer the question which functions the two UPF3 paralogs carry out in human cells. Since the initial description of mammalian UPF3A and UPF3B (Lykke-Andersen *et al*., 2000; Serin *et al*., 2001), many researchers have been engaged in determining the function and work distribution of these two proteins in NMD. It became clear relatively early that UPF3A and UPF3B can both interact with UPF2 as well as the EJC and activate NMD (Gehring *et al*., 2003; Kim *et al*., 2001; Lykke-Andersen *et al*., 2000; Serin *et al*., 2001). But significant differences in these interactions and the amounts of UPF3A and UPF3B proteins were also found, leading to the hypothesis that UPF3B is the central player and UPF3A more its backup (Chan *et al*., 2009; Kunz *et al*., 2006). The investigations were further intensified upon the discovery that mutations in the human UPF3B gene lead to various forms of mental disorder (Nguyen *et al*., 2014). Disease severity seems to be dictated by the amount of UPF3A present, suggesting again a compensatory mechanism and redundant functions of UPF3B and UPF3A (Nguyen *et al*., 2012).

Later, it was reported that in mouse cells UPF3B is an NMD activator and UPF3A is an NMD inhibitor (Shum *et al*., 2016). Accordingly, removal of UPF3A resulted in enhanced NMD, whereas removal of UPF3B inhibited NMD. Recently, another function of UPF3B was discovered, namely that it is involved in different phases of translation termination (Neu-Yilik *et al*., 2017). UPF3B not only interacts with release factors, but also slows down translation termination and promotes dissociation of post-termination ribosomal complexes. Whether these functions of UPF3B in translation termination are related to its function in NMD has not been clarified to date.

Although our results cannot fully answer this last question, we have more or less definite answers for the functions of the two proteins UPF3A and UPF3B in NMD. All our data support the notion that the presence of UPF3B or UPF3A is sufficient to maintain NMD activity in human cells. First, we see at most a weak inhibition of NMD in our HEK293 UPF3B KO cells. However, this does not necessarily mean that all cell types display full NMD activity after a KO of UPF3B. Instead, it is conceivable that HEK293 cells are just particularly robust against the UPF3B depletion or particularly efficient at the compensatory upregulation of UPF3A. Likewise, we see no NMD inhibition in cells overexpressing UPF3A or NMD “boosting” in UPF3A KO cells. Only when we deplete in UPF3A or UPF3B KO cell lines the respective other protein by RNAi or genomic KO, NMD efficiency substantially decreases. It should be noted that the effects of a KO on NMD activity are typically stronger than those of a KD, which is consistent with our earlier observations (Boehm *et al*., 2021; Gerbracht *et al*., 2020). Although our data clearly argue against an NMD-inhibitory function of UPF3A, experimental differences exist between our work and the work describing the NMD inhibition by UPF3A. While we have used human HEK293 cells, the results of Shum et al. were obtained in P19 mouse cells and mouse embryonic fibroblasts (MEFs). Both, the different organisms and the different types of cells could have influenced the results. Interestingly, in a parallel manuscript, Yi et al. find that mouse UPF3A can rescue NMD in human HCT116 UPF3A-UPF3B dKO cells (Yi *et al*., 2021). Although these results were obtained in a heterologous context, mouse UPF3A does not appear to be a general NMD inhibitor.

The different KO cells that we have generated in the course of this project enabled us to conduct experiments that went beyond investigating UPF3-dependent NMD substrates in human cells. Specifically, we were able to study the composition of NMD complexes without UPF3B or both UPF3 proteins and to carry out rescue experiments with different UPF3A and UPF3B protein variants. Not entirely unexpected, we observed that in the absence of UPF3A and UPF3B, the interaction between UPF2 and the EJC is lost. This bridging by UPF3 between UPF2-containing NMD complexes and the EJC was previously considered to be essential for NMD. However, two observations argue against UPF3 being mainly a bridging protein. First, UPF3B mutants that cannot interact with either the EJC or UPF2 fully rescue NMD. Only when both interaction sites were mutated, UPF3 lost its NMD function. This indicates that the interaction with one of the two interaction partners is sufficient to maintain NMD. Second, we observed that not only in UPF3 double-KO cells, but also in UPF3B KO cells, the bridge between UPF2 and the EJC was lost. Although quite surprising at first glance, this is in good agreement with previous results showing that the interaction between the EJC and UPF3A is substantially weaker than that between the EJC and UPF3B (Kunz *et al*., 2006). Indeed, earlier structural data also argue against a bridging function of UPF3. In the cryo-EM structure of an EJC-UPF3-UPF2-UPF1 complex, UPF1 did not face towards a possible terminating ribosome in the 5’ direction, but instead in 3’ direction (Melero *et al*., 2012). Therefore, one could conclude that the interactions between all these proteins do not take place at a single time point during NMD.

This raises the question of what UPF3 function is essential for NMD, if it is not its bridging function? Our rescue experiments showed that the middle domain of UPF3B cooperates with the UPF2- and the EJC interaction sites, i.e., its deletion in combination with one other mutation inactivates UPF3B. The middle domain has been described to mediate the interaction of UPF3B with release factor 3 (RF3), but this interaction has not been demonstrated for UPF3A (Neu-Yilik *et al*., 2017). So, if UPF3A cannot interact with RF3 and also binds weaker to the EJC as noted above, it should be functionally inactive. However, we see no obvious difference between UPF3B and UPF3A in the rescue experiments. These and other observations can, in our view, only be explained with more complex models, which must also consider non-linear relationships and potential auxiliary functions of certain regions of UPF3 (Fig 8).

**Figure 8.**
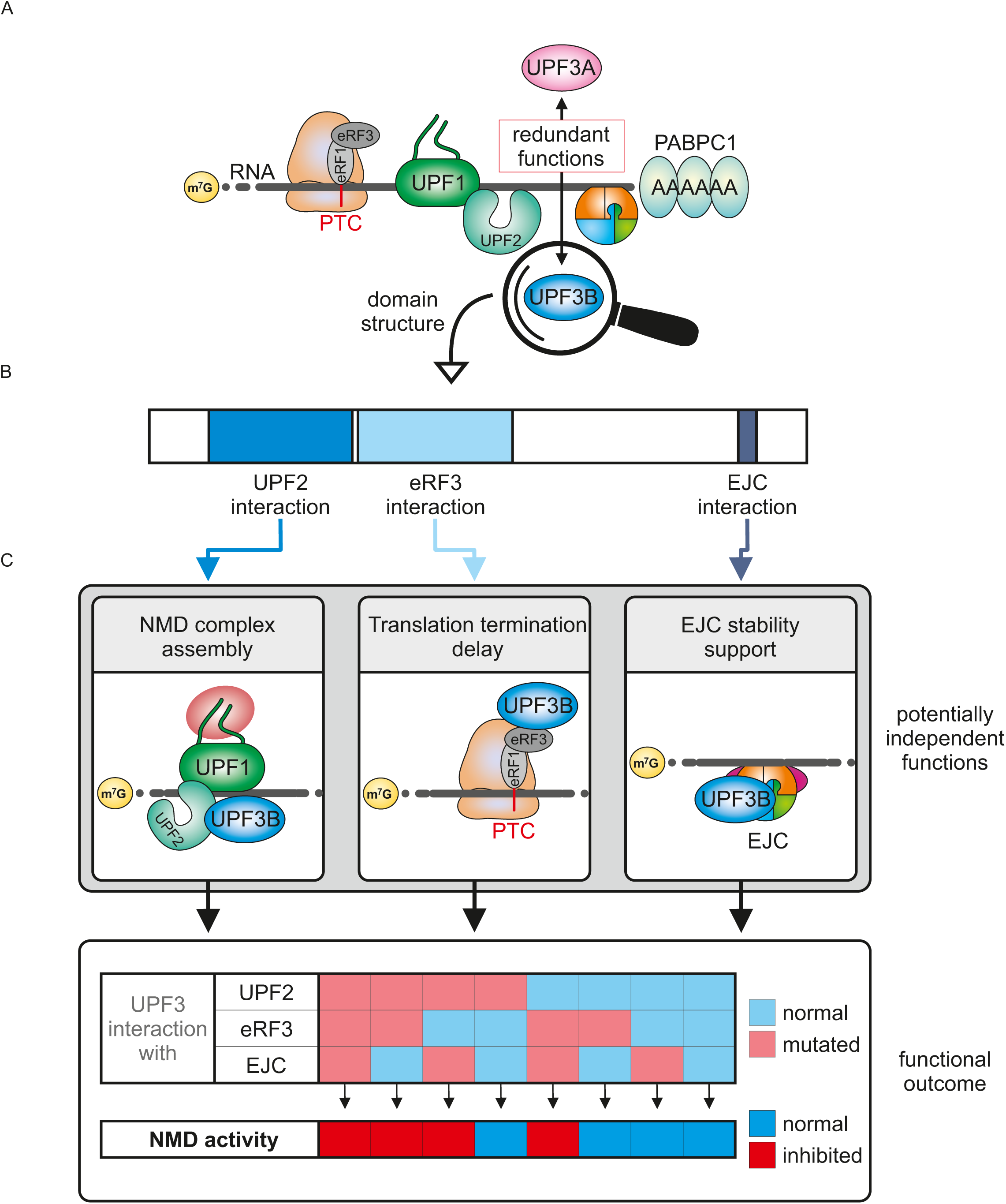
Model for potentially independent functions of UPF3 in NMD. **A** Both UPF3 paralogs UPF3Aand UPF3B can elicit NMD and trigger mRNAdegradation. We propose a new model where UPF3B exerts multiple functions at different timepoints during NMD. **B** Schematic overview of UPF3B domain structure and the postulated functions. **C** Via the RRM-like domain in the N-terminus of the protein it interacts with UPF2, which might be important for the assembly of an NMD-inducing complex. The middle domain was ascribed to be responsible for the interaction with eRF3 and potential translation termination delay. The interaction with the EJC via the EBM could stabilize the complex by preventing its interaction with the disassembly factor PYM1. Only if two out of the three functions are ensured, NMD can pursue and degrade the targeted mRNA.

Since the interactions of UPF3 are essential only in combination with each other, we propose that UPF3A and UPF3B exert multiple functions at different time points of NMD and in association with different complexes. We only consider here the previously described interactions of UPF3 with UPF2, the EJC and the release factor 3 (Fig 8B). While it was previously described that UPF3B interacts better than UPF3A with UPF2 (Chan *et al*., 2009), we find both proteins in the FLAG-UPF2 IP. The amount of UPF3A does not seem to increase when UPF3B is depleted, which could be due to the overexpression of UPF2 in our experimental system. The interaction of UPF3 and UPF2 is also conserved in yeast and is thus likely to have functional significance. However, a UPF3B mutant in which binding to UPF2 was inactivated rescues NMD better than the UPF3B WT.

The interaction of the middle domain of UPF3B with RF3 has only recently been described (Neu-Yilik *et al*., 2017). Again, we find that removing only the middle domain does not substantially inhibit NMD. In combination with an inactivation of the UPF2 binding, the deletion of the middle domain leads to a complete inhibition of the NMD in the rescue assay. This could also happen when exon 4 of UPF3B or UPF3A is removed, which is located at the junction between the UPF2 binding domain and the middle domain. Therefore, UPF3A Δexon4 would be a naturally occurring, NMD-inactive variant of the UPF3 proteins.

How might the different regions and domains communicate with each other and regulate the function of UPF3 in NMD? The function of the middle domain in relation to NMD has not yet been investigated. It is conceivable that UPF3B plays a minor role in translation termination and that the events that trigger NMD can also occur without the middle domain - potentially with a delay (Figure 8c). Therefore, the deletion of the middle domain could be tolerated in isolation but would become fatal in combination with other mutations that impair additional functions. With regard to the EBM, we propose that its binding stabilizes the EJC, for example by preventing the interaction of the EJC with PYM1 (Fig 8C). PYM1 is a known EJC disassembly factor and binds to the EJC at a surface area that overlaps with the EBM binding site (Bono *et al*., 2004; Buchwald *et al*., 2010; Gehring *et al*., 2009). We suggest that in cells rescued with the ΔEBM mutant EJCs are more readily dissociated from the mRNA in the cytoplasm. The concomitant loss of UPF3B’s termination function (Δmiddle domain, ΔEBM) would have a dramatic effect on NMD efficiency, because NMD would be initiated too slowly. Likewise, the interaction of UPF3 with UPF2 might be important for the assembly of an NMD- inducing complex (Fig 8C) that needs to be timed with translation termination and that leads to NMD activation only in the presence of the EJC. Although these suggestions may not accurately reflect the molecular events during NMD, they illustrate possible functions of the domains of UPF3, particularly in relation to their NMD-inactive combinations (Fig 8). Overall, our observations fit well with a "synthetic lethal" model in which inactivation of any two domains together disrupts UPF3 activity.

One factor whose function needs to be examined in more detail in the context of UPF3 is CASC3. Our mass spectrometry analysis shows that CASC3 immunoprecipitates very well with UPF2 in wildtype cells. CASC3 still partially precipitates with UPF2 in UPF3B KO cells, although the interaction of the other EJC factors is reduced to background levels. In previous work, we observed that UPF3B interacts less well with the EJC when CASC3 is knocked out (Gerbracht *et al*., 2020). This indicates that an interaction between CASC3 and the UPF3 proteins exists that is not well understood so far. What kind of interaction this is and what function it has will be interesting to address in future experiments.

Our own work and the work of Yi et al. have re-examined the functions of the human UPF3 paralogues UPF3A and UPF3B (Yi *et al*., 2021). Together, the studies confirmed some previous findings and disconfirmed others, thereby successfully (re-)defining the role of UPF3 proteins in human cells. A few questions remain unanswered and need to be addressed in the future, for example how exactly the middle domain supports NMD. As described above, our results have implications for the understanding of the NMD mechanism, as they are incompatible with, and thus exclude, certain models of NMD. In addition, they may also help to better understand the link between UPF3B and intellectual disability and which domains of UPF3A modulate the severity of the disease and may therefore be potential targets for therapy.

## Materials and Methods

### Cell Culture

Flp-In T-Rex 293 (human, female, embryonic kidney, epithelial; Thermo Fisher Scientific, RRID:CVCL_U427) cells were cultured in high-glucose, GlutaMAX DMEM (Gibco) supplemented with 9% fetal bovine serum (Gibco) and 1x Penicillin Streptomycin (Gibco). The cells were cultivated at 37°C and 5% CO2 in a humidified incubator. The generation of knockout and stable cell lines is described below and all cell lines are summarized in Dataset EV6.

### siRNA-mediated knockdowns

For reverse transfection the cells were seeded at a density of 2.5 x 10^5^ cells per well. The transfection solution contained 2.5 µl Lipofectamine RNAiMAX and 60 pmol of the respective siRNAs. For the UPF3A and UPF3B knockdowns 30 pmol of both belonging siRNAs were used. In preparation for mass spectrometry, 2.5 x 10^6^ were reverse transfected in 10 cm plates using 6.25 µl Lipofectamine RNAiMAX and 150-200 pmol siRNA (or half of it for each of the two siRNAs for UPF3A or UPF3B). All siRNAs used in this study are listed in Dataset EV6.

### Plasmid transfection

For each stable transfection 2.5-3.0 x 10^5^ cells were seeded one day prior transfection in 6-wells. To express the N-terminally FLAG-tagged protein constructs and reporter mRNAs for northern blotting, they were stably integrated using the PiggyBac (PB) Transposon system with the cumate-inducible PB-CuO-MCS-BGH-EF1-CymR-Puro vector. This vector was modified from the original vector (PB-CuO-MCS-IRES-GFP-EF1α-CymR-Puro (System Biosciences)) by replacing the IRES-GFP cassette with a BGH polyA signal. Per well 1.0 μg of the respective PB vector and 0.8 μg PB Transposase were transfected using a calcium phosphate-based system with BES buffered saline (BBS). Additionally, 0.5 μg of pCI-maxGFP was transfected as a visual feedback for transfection efficiency. 48 h later, the cells were pooled in 10 cm plates and selected for positive cells by incubation in media containing 2 μg/ml puromycin for a week. To induce expression of the constructs, 30 μg/ml cumate was added and the cells were harvested after 72h for continuing experiments.

The mRNA reporter constructs β-globin WT and β-globin PTC are available on Addgene (IDs 108375-108376). All vectors used in this study are listed in Dataset EV6.

### Generation of knockout cells using CRISPR-Cas9

The knockouts were performed using the Alt-R CRISPR-Cas9 system (Integrated DNA Technologies) and reverse transfection of a Cas9:guideRNA ribonucleoprotein complex using Lipofectamine RNAiMAX (Thermo Fisher Scientific) according to the manufacturer’s protocol. The crRNA sequence (Integrated DNA Technologies) to target UPF3B was /AltR1/rArGrArUrArArGrCrArGrGrArUrCrGrCrArArCrArGrUrUrUrUrArGrArGrCrUrArUrGrCrU/ AltR2/. For UPF3A the crRNA sequences were /AltR1/rCrCrGrCrArArCrCrGrGrArGrGrArC rGrArArGrUrGrUrUrUrUrArGrArGrCrUrArUrGrCrU/AltR2/ for clone 1 and /AltR1/rGrCrGrGr UrGrGrArArCrUrGrCrArCrUrUrCrUrArGrUrUrUrUrArGrArGrCrUrArUrGrCrU/AltR2/ for clone Reverse transfection was performed on 1.5×10^5^ cells per crRNA in 12-well plates. 48 h after transfection the cells were trypsinized, counted and seeded at a mean density of a single cell per well in 96-well plates. Cell colonies originating from a single clone were then screened via Western blot and genome editing of UPF3A and UPF3B was analyzed on the genomic level via DNA extraction and Sanger sequencing. Alterations on the transcript level were analyzed via RNA extraction followed by reverse transcription and Sanger sequencing.

### DNA and RNA extraction

Genomic DNA extraction using QuickExtract DNA Extraction Solution (Lucigen) was performed according to manufacturer’s instruction. For RNA extraction cells were harvested with 1 ml RNAsolv reagent (Omega Bio-Tek) per 6 well and RNA was isolated according to manufacturer’s instruction, with the following changes: instead of 200 μl chloroform, 150 μl 1-Bromo-3-chloropropane (Sigma-Aldrich) was added to the RNAsolv. Additionally, in the last step the RNA pellet was dissolved in 20 μl RNase-free water by incubating for 10 min on a shaking 65 °C heat block.

### Western blotting

SDS-polyacrylamide gel electrophoresis and immunoblot analysis were performed using protein samples harvested with RIPA buffer (50 mM Tris/HCl pH 8.0, 0.1% SDS, 150 mM NaCl, 1% IGEPAL, 0.5% deoxycholate) or samples eluted from Anti-FLAG M2 magnetic beads. For protein quantification, the Pierce Detergent Compatible Bradford Assay Reagent (Thermo Fisher Scientific) was used. All antibodies used in this study are listed in Dataset EV6. Detection was performed with Western Lightning Plus-ECL (PerkinElmer) or ECL Select Western Blotting Detection Reagent (Amersham) and the Vilber Fusion FX6 Edge imaging system (Vilber Lourmat).

### Semi-quantitative and quantitative reverse transcriptase (RT)-PCR

Reverse transcription was performed with 1-4 µg of total RNA in a 20 µl reaction volume with 10 µM VNN-(dT)_20_ primer and the GoScript Reverse Transcriptase (Promega). For the semi-quantitative end-point PCRs the MyTaq Red Mix (Bioline) was used. Quantitative RT-PCRs were performed with the GoTaq qPCR Master Mix (Promega), 2% of cDNA per reaction, and the CFX96 Touch Real-Time PCR Detection System (Bio-Rad). Each biological replicate was repeated in technical triplicates and the average Ct (threshold cycle) value was measured. When isoform switches were measured, values for NMD sensitive isoforms were normalized to the canonical isoforms to calculate ΔCt. For differentially expressed genes, the housekeeping gene C1orf43 values were subtracted from the target value to receive the ΔCt. To calculate the mean log2 fold changes three biologically independent experiments were used. The log2 fold changes are visualized as single data points and mean. All primers used in this study are listed in Dataset EV6.

### RNA-sequencing and computational analyses

Four different RNA-seq experiments were performed: 1) unaltered Flp-In T-REx 293 wild type (WT) cells and WT cells overexpressing the UPF3A WT construct via the PB-Transposase system. 2) Flp-In T-REx 293 wild type (WT) cells transfected with Luciferase siRNA and the UPF3A KO clones 14 and 20 treated with either Luciferase or UPF3B siRNAs. 3) The control Flp-In T-REx 293 wild type (WT) cells with a Luciferase KD and the UPF3B KO clone 90 transfected with Luciferase or UPF3A siRNAs. 4) WT cells transfected with Luciferase siRNA and the two UPF3 double KO cell lines 1 and 2 transfected with either Luciferase or UPF3B siRNAs. RNA was purified using peqGOLD TriFast (VWR Peqlab; for UPF3B KO samples) or the Direct-zol RNA MiniPrep kit including the recommended DNase I treatment (Zymo Research; all other samples) according to manufacturer’s instructions. Three biological replicates were analyzed for each sample.

The Lexogen SIRV Set3 Spike-In Control Mix (SKU: 051.0x; for UPF3B KO samples) or ERCC RNA Spike-In Mix (for all other samples) that provides a set of external RNA controls was added to the total RNA to enable performance assessment. The Spike-Ins were used for quality control purposes, but not used for the final analysis of DGE, DTU or AS.

Using the Illumina TruSeq Stranded Total RNA kit library preparation was accomplished. This includes removing ribosomal RNA via biotinylated target-specific oligos combined with Ribo-Zero gold rRNA removal beads from 1 μg total RNA input. Cytoplasmic and mitochondrial rRNA gets depleted by the Ribo-Zero Human/Mouse/Rat kit. After a purification step, the RNA gets cleaved and fragmented. These fragments are then reverse transcribed into first strand cDNA using reverse transcriptase and random primers. In the next step, using DNA Polymerase I and RNase H second strand cDNA synthesis is performed. The resulting cDNA fragments then have the extension of a single’A’ base and adapter ligation. To create the final cDNA library the products are purified and enriched with PCR. Next library validation and quantification (Agilent tape station) are performed, followed by pooling of equimolar amounts of library. The pool itself was then quantified using the Peqlab KAPA Library Quantification Kit and the Applied Biosystems 7900HT Sequence Detection System and sequenced on an Illumina HiSeq4000 sequencing instrument with an PE75 protocol (UPF3B KO samples) or Illumina NovaSeq6000 sequencing instrument with an PE100 protocol (all other samples).

Reads were aligned against the human genome (version 38, GENCODE release 33 transcript annotations (Frankish *et al*., 2019) supplemented with SIRVomeERCCome annotations from Lexogen; obtained from https://www.lexogen.com/sirvs/download/) using the STAR read aligner (version 2.7.3a) (Dobin *et al*., 2013). Transcript abundance estimates were computed with Salmon (version 1.3.0) (Patro *et al*., 2017) with a decoy-aware transcriptome. After the import of transcript abundances, differential gene expression analysis was performed with the DESeq2(Love *et al*., 2014) R package (version 1.28.1) with the significance thresholds |log2FoldChange|> 1 and adjusted p-value (padj) < 0.05. Differential splicing was detected with LeafCutter (version 0.2.9) (Li *et al*., 2018) with the significance thresholds |deltapsi| > 0.1 and adjusted p-value (p.adjust) < 0.05. Differential transcript usage was computed with IsoformSwitchAnalyzeR (version 1.10.0) and the DEXSeq method (Anders *et al*., 2012; Ritchie *et al*., 2015; Robinson & Oshlack, 2010; Soneson *et al*., 2015; Vitting-Seerup & Sandelin, 2017, 2019). Significance thresholds were |dIF| > 0.1 and adjusted p-value (isoform_switch_q_value) < 0.05.

PTC status of transcript isoforms with annotated open reading frame was determined by IsoformSwitchAnalyzeR using the 50 nucleotide (nt) rule of NMD (Huber *et al*., 2015; Vitting-Seerup *et al*., 2014; Vitting-Seerup & Sandelin, 2017; Weischenfeldt *et al*., 2012). Isoforms with no annotated open reading frame in GENCODE were designated “NA” in the PTC analysis.

All scripts and parameters for the RNA-Seq analysis are available at GitHub [https://github.com/boehmv/UPF3]. Overlaps of data sets were represented via nVenn(Perez-Silva *et al*., 2018) or the ComplexHeatmap package (version 2.6.2)(Gu *et al*., 2016). Integrative Genomics Viewer (IGV) (version 2.8.12)(Robinson *et al*., 2011) snapshots were generated from mapped reads (BAM files) converted to binary tiled data (tdf), using Alfred(Rausch *et al*., 2019) with resolution set to 1 and IGVtools.

### SILAC and mass spectrometry

HEK293 WT cells and the UPF3 dKO clone 2 expressing either FLAG-tagged GST or UPF2 were labeled by culturing them for at least 5 passages in DMEM for SILAC medium (Thermo Fisher Scientific) supplemented with 9% FBS (Silantes), 1% Penicillin-Streptomycin and the labeled amino acids Lysin and Arginine at final concentrations of 0.798 mmol/L and 0.393 mmol/L, respectively. The three conditions were “light” (unlabeled Lys/ Arg), “medium” (Lys4/ Arg6) and “heavy” (Lys8/ Arg10). Unlabeled proline was added in all conditions to prevent enzymatic Arginine-to-Proline conversion.

### Experimental setup for SILAC with FLAG-tagged UPF2

Expression of FLAG-GST and FLAG-UPF2 was induced for 72 h with 1x cumate. The cells lysed in 250 – 400 μl Buffer E with 1 μg/ml RNase and sonicated using the Bandelin Sonopuls mini20 with 15x 1s pulses at 50% amplitude with a 2.5 mm tip. Protein concentrations were measured using the Bradford assay and protein samples containing 1.6-1.7 mg/ml total protein were diluted. 600 μl of these samples were incubated with 30 μl Anti-FLAG M2 magnetic beads (Sigma) for 2 h on an overhead shaker at 4 °C. The beads were then washed three times for 5 min with mild EJC-Buffer before eluting twice with 22 μl of a 200 μg/ml dilution of FLAG-peptides (Sigma) in 1x TBS for 10 min at RT and 200 rpm each elution step. Another elution with 1x SDS loading buffer was performed to analyze pull down efficiency via Western blot. The FLAG-peptide eluates were then mixed as followed: 7 μl of both light conditions, 14 μl medium and 14 μl heavy. 1 volume of SP3 (10% SDS in PBS) was added and the samples were reduced with 5 mM DTT and alkylated with 40 mM CAA.

Tryptic protein digestion was achieved by following a modified version of the single pot solid phase-enhanced sample preparation (SP3) (Hughes *et al*., 2014). In brief, paramagnetic Sera-Mag speed beads (Thermo Fisher Scientific) were added to the reduced and alkylated protein samples and then mixed 1:1 with 100% acetonitrile (ACN). Protein-beads-complexes form during the 8 min incubation step, followed by capture using an in-house build magnetic rack. After two washing steps with 70% EtOH, the samples were washed once with 100% ACN. Then they were air-dried, resuspended in 5 μl 50 mM Triethylamonium bicarbonate supplemented with trypsin and LysC in an enzyme:substrate ratio of 1:50 and incubated for 16 h at 37°C. The next day the beads were again resuspended in 200 μl ACN and after 8 min incubation placed on the magnetic rack. Tryptic peptides were washed with 100% ACN and air-dried before dissolved in 4% DMSO and transfer into 96-well PCR tubes. The last step was the acidification with 1 μl of 10% formic acid, then the samples were ready for mass spec analysis.

Proteomics analysis was performed by the proteomics core facility at CECAD via data-dependent acquisition using an Easy nLC1200 ultra high-performance liquid chromatography (UHPLC) system connected via nano electrospray ionization to a Q Exactive Plus instrument (all Thermo Scientific) running in DDA Top10 mode. Based on their hydrophobicity the tryptic peptides were separated using a chromatographic gradient of 60 min with a binary system of buffer A (0.1% formic acid) and buffer B (80% ACN, 0.1% formic acid) with a total flow of 250 nl/min. For the separation in-house made analytical columns (length: 50 cm, inner diameter: 75 μm) containing 2.7 μm C18 Poroshell EC120 beads (Agilent) that were heated to 50 °C in a column oven (Sonation) were used. Over a time period of 41 min Buffer B was linearly increased from 3% to 27% and then more rapidly up to 50% in 8 min. Finally, buffer B was increased to 95% within 1 min followed by 10 min at 95% to wash the analytical column. Full MS spectra (300-1,750 m/z) were accomplished with a resolution of 70,000, a maximum injection time of 20 ms and an AGC target of 3e6. In each full MS spectrum, the top 10 most abundant ions were selected for HCD fragmentation (NCE:27) with a quadrupole isolation width of 1.8 m/z and 10 s dynamic exclusion. The MS/MS spectra were then measured with a 35,000 resolution, an injection time of maximum 110 ms and an AGC target of 5e5.

The MS RAW files were then analyzed with MaxQuant suite (version 1.5.3.8) on standard settings with the before mentioned SILAC labels (Cox & Mann, 2008). By matching against the human UniProt database the peptides were then identified using the Andromeda scoring algorithm (Cox *et al*., 2011). Carbamidomethylation of cysteine was defined as a fixed modification, while methionine oxidation and N-terminal acetylation were variable modifications. The digestion protein was Trypsin/P. A false discovery rate (FDR) < 0.01 was used to identify peptide-spectrum matches and to quantify the proteins. Data processing and statistical analysis was performed in the Perseus software (version 1.6.1.1) (Tyanova *et al*., 2016). Using the One-sample t-test the significantly changed proteins were identified (H0 = 0, fudge factor S0 = 0.2). Visualization was performed with RStudio (version 1.2.5033).

### Label-free quantitative mass spectrometry

Twenty-four hours before expression of the FLAG-tagged constructs, the HEK293 WT cells were treated with Luciferase siRNA and the UPF3B KO clone 90 and UPF3 dKO clone 1 cells were treated with siRNAs targeting residual UPF3B. The expression of either FLAG-GST or FLAG-UPF2 in WT cells and FLAG-UPF2 in the clones 90 and 1 was induced for 48 h with 1x. Lysis and sample preparation were performed as described above. MS analysis was performed as described above with a slightly adjusted gradient as followed: 3 – 30% B in 41 min, 30 – 50% B in 8 min, 50-95% B in 1 min, followed by 10 min washing at 95%. LFQ values were calculated using the MaxLFQ algorithm (Cox *et al*., 2014) in MaxQuant. Significantly changed proteins were identified by two-sample *t-*testing (fudge factor S0 = 0.2).

### Northern Blotting

The cells were harvested in RNAsolv reagent and total RNA extraction was performed as described above. 3.0 µg total RNA were resolved on a 1% agarose/0.4 M formaldehyde gel using the tricine/triethanolamine buffer system (Mansour & Pestov, 2013). Next a transfer on a nylon membrane (Roth) in 10x SSC followed. The blot was incubated overnight at 65°C in Church buffer containing [α-32P]-GTP body-labeled RNA-probes for mRNA reporter detection (Voigt *et al*., 2019). Ethidium bromide stained 28S and 18S rRNA served as loading controls. RNA signal detected with the Typhoon FLA 7000 (GE Healthcare) was quantified in a semi-automated manner using the ImageQuant TL 1D software with a rolling-ball background correction. EtBr-stained rRNA bands were quantified with the Image Lab 6.0.1 software (Bio-Rad). Signal intensities were normalized to the internal control (rRNA) before calculation of mean values. The control condition was set to unity (TPI WT for reporter assays), quantification results are shown as data points and mean.

### Protein conservation

UPF3A (UniProt ID: Q9H1J1-1) and UPF3B (UniProt ID: Q9BZI7-2) protein sequences were aligned using Clustal Omega (https://www.ebi.ac.uk/Tools/msa/clustalo/) (Goujon *et al*., 2010; Sievers *et al*., 2011), viewed using Jalview (Waterhouse *et al*., 2009), the conservation score extracted and used for visualization.

### Data Presentation

Quantifications and calculations for other experiments were performed - if not indicated otherwise - with Microsoft Excel (version 1808) or R (version 4.0.4) and all plots were generated using IGV (version 2.8.12), GraphPad Prism 5, ggplot2 (version 3.3.3) or ComplexHeatmap (version 2.6.2) (Gu *et al*., 2016).

### Data Availability

The datasets and computer code produced in this study are available in the following databases:

• RNA-Seq data for UPF3B KO samples: ArrayExpress E-MTAB-10711 (https://www.ebi.ac.uk/arrayexpress/experiments/E-MTAB-10711)
• RNA-Seq data for UPF3 dKO samples: ArrayExpress E-MTAB-10716 (https://www.ebi.ac.uk/arrayexpress/experiments/E-MTAB-10716)
• RNA-Seq data for UPF3A KO/OE samples: ArrayExpress E-MTAB-10718 (https://www.ebi.ac.uk/arrayexpress/experiments/E-MTAB-10718)
• Mass spectrometry proteomics data: PRIDE PXD027120 (https://www.ebi.ac.uk/pride/archive/projects/PXD027120)
• Codes used in this study: GitHub (https://github.com/boehmv/UPF3)

All relevant data supporting the key findings of this study are available within the article and its Expanded View files or from the corresponding author upon reasonable request.

## Acknowledgements

We thank members of the Gehring lab for discussions and reading of the manuscript. We also thank Marek Franitza and Christian Becker (Cologne Center for Genomics, CCG) for preparing the sequencing libraries and operating the sequencer. This work was supported by grants from the Deutsche Forschungsgemeinschaft to C.D. (DI 1501/8-1, DI1501/8-2) and N.H.G (GE 2014/6-2 and GE 2014/10-1) and by the Center for Molecular Medicine Cologne (CMMC, Project C 07; to N.H.G.). V.B. was funded under the Institutional Strategy of the University of Cologne within the German Excellence Initiative. N.H.G. acknowledges funding by a Heisenberg professorship (GE 2014/7-1 and GE 2014/13-1) from the Deutsche Forschungsgemeinschaft. C.D was kindly supported by the Klaus Tschira Stiftung gGmbH (00.219.2013). This work was supported by the DFG Research Infrastructure as part of the Next Generation Sequencing Competence Network (project 423957469). NGS analyses were carried out at the production site WGGC Cologne.

## Author contributions

Conceptualization: Niels H. Gehring, Volker Boehm and Damaris Wallmeroth;

Methodology: Volker Boehm, Damaris Wallmeroth, Niels H. Gehring;

Software: Volker Boehm;

Investigation: Damaris Wallmeroth, Volker Boehm, and Jan-Wilm Lackmann;

Resources and Data Curation: Volker Boehm, Janine Altmüller and Jan-Wilm Lackmann;

Writing – Original Draft, Review & Editing: Damaris Wallmeroth, Volker Boehm, Niels H. Gehring;

Visualization: Volker Boehm and Damaris Wallmeroth;

Supervision: Niels H. Gehring and Volker Boehm;

Funding Acquisition: Niels H. Gehring and Christoph Dieterich;

