## Supplemental Figures for "UPF3A and UPF3B are redundant and modular activators of nonsense-mediated mRNA decay in human cells"

### Expanded View Figures

**This PDF file includes:**

Figure EV1 to EV6

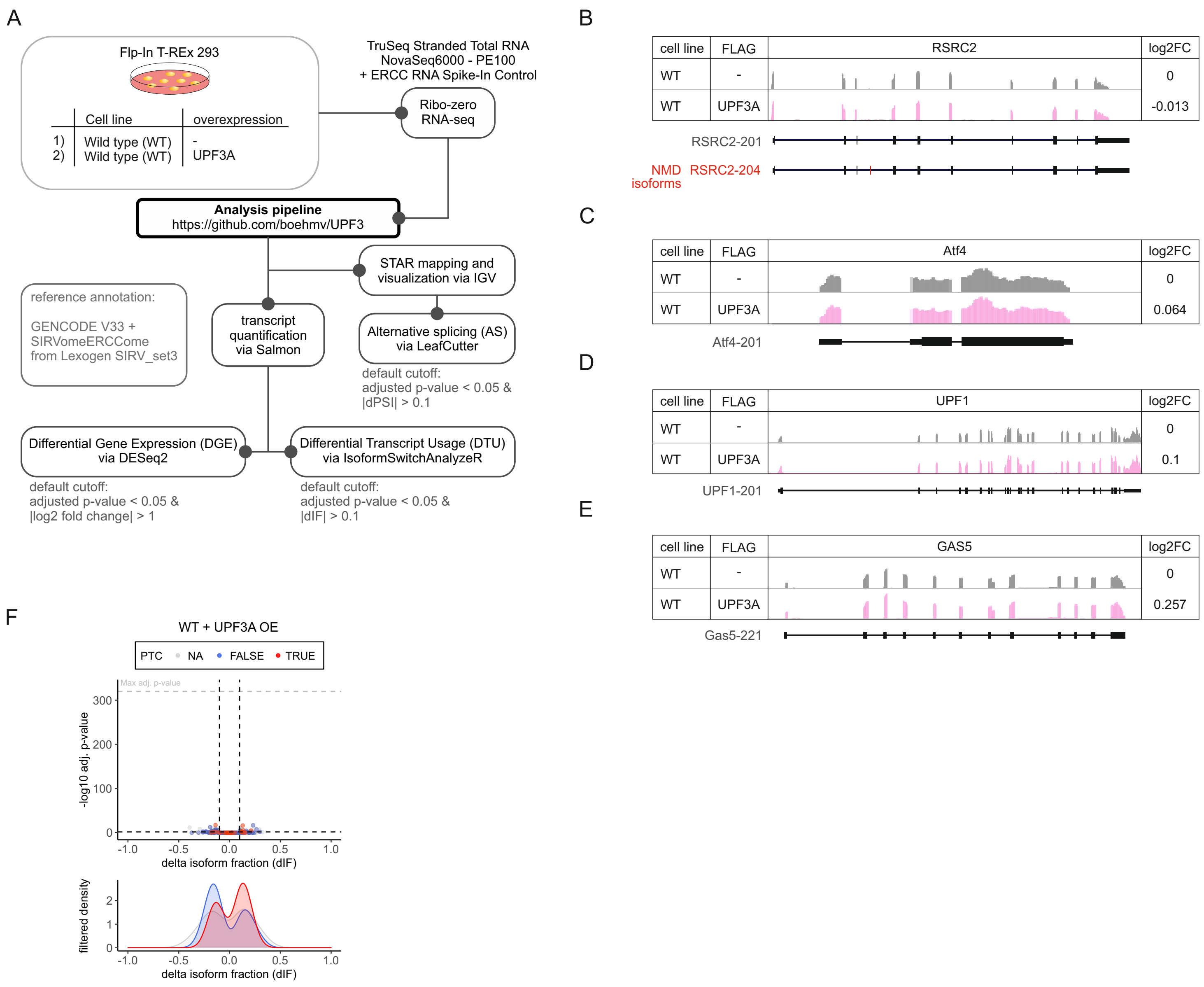


**Figure EV1 - UPF3A overexpression does not cause upregulation of NMD-targets**

**A** Schematic overview of the analysis pipeline.

**B-E** Read coverage of indicated genes from WT RNA-seq data with or without induced UPF3A overexpression shown as Integrative Genomics Viewer (IGV) snapshot. Differential gene expression (from DESeq2) is indicated as log2 fold change (log2FC) on the right. Schematic representation of the protein coding transcript below.

**F** Volcano plot showing the differential transcript usage (via IsoformSwitchAnalyzeR) in RNA-Seq data of WT cells overexpressing UPF3A. Isoforms containing GENCODE (release 33) annotated PTC (red, TRUE), regular stop codons (blue, FALSE) or having no annotated open reading frame (grey, NA) are indicated. The change in isoform fraction (dIF) is plotted against the -log10 adjusted p-value (padj). Density plots show the distribution of filtered isoforms in respect to the dIF, cutoffs were |dIF| > 0.1 and adj. p-value < 0.05. P-values were calculated by IsoformSwitchAnalyzeR using a DEXSeq-based test and corrected for multiple testing using the Benjamini-Hochberg method. OE = overexpression.

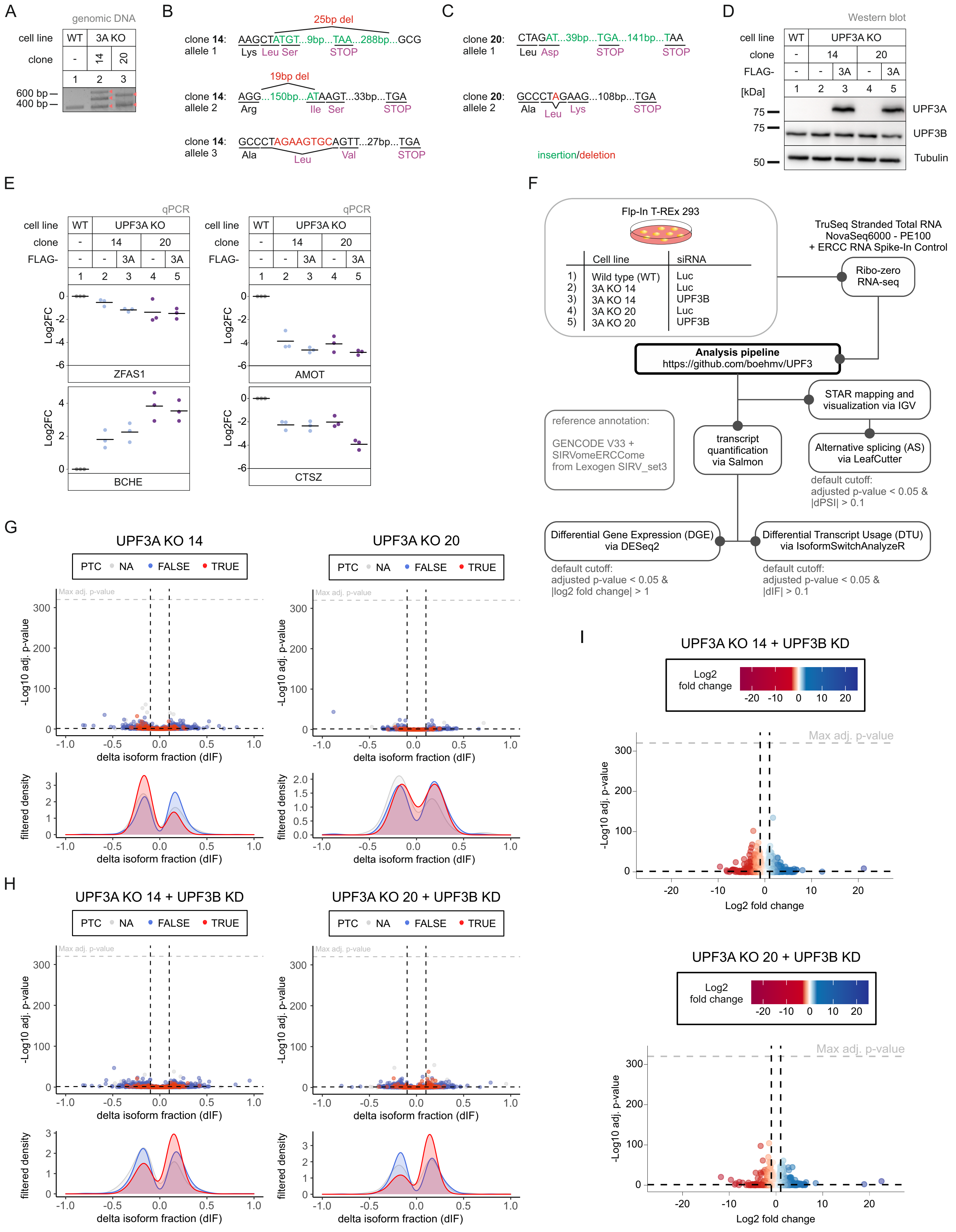


**Figure EV2 - Light NMD inhibition after UPF3B KD in UPF3A KO cells**

**A** PCR-amplification of targeted genomic locus of UPF3A for Sanger sequencing analysis.

**B,C** The targeted exon region and anticipated PTC location following insertions (green) or deletions (red) are indicated for detected alleles of UPF3A in clone 14 (B) and 20 (C).

**D** Western blot analysis of WT and UPF3A KO cells (clones 14 and 20) with or without expression of FLAG-tagged UPF3A rescue construct. UPF3A and UPF3B protein levels were detected, Tubulin serves as control.

**E** Quantitative RT-PCR of the samples from (D). Expression of four targets with significant DGE in both UPF3A KO clones was normalized to C1orf43 reference. Data points and means are plotted as log2 fold change (log2FC) (n=3).

**F** Schematic overview of the analysis pipeline.

**G,H** Volcano plots showing the differential transcript usage (via IsoformSwitchAnalyzeR) in various RNA-Seq data. Isoforms containing GENCODE (release 33) annotated PTC (red, TRUE), regular stop codons (blue, FALSE) or having no annotated open reading frame (grey, NA) are indicated. The change in isoform fraction (dIF) is plotted against the -log10 adjusted p-value (adj. p-value). Density plots show the distribution of filtered isoforms in respect to the dIF, cutoffs were |dIF| > 0.1 and adj.p-value < 0.05. P-values were calculated by IsoformSwitchAnalyzeR using a DEXSeq-based test and corrected for multiple testing using the Benjamini-Hochberg method.

**I** Volcano plots showing the differential gene expression analyses from the indicated RNA-Seq datasets. The log2 fold change is plotted against the -log10 adjusted p-value (padj). P-values were calculated by DESeq2 using a two-sided Wald test and corrected for multiple testing using the Benjamini-Hochberg method.

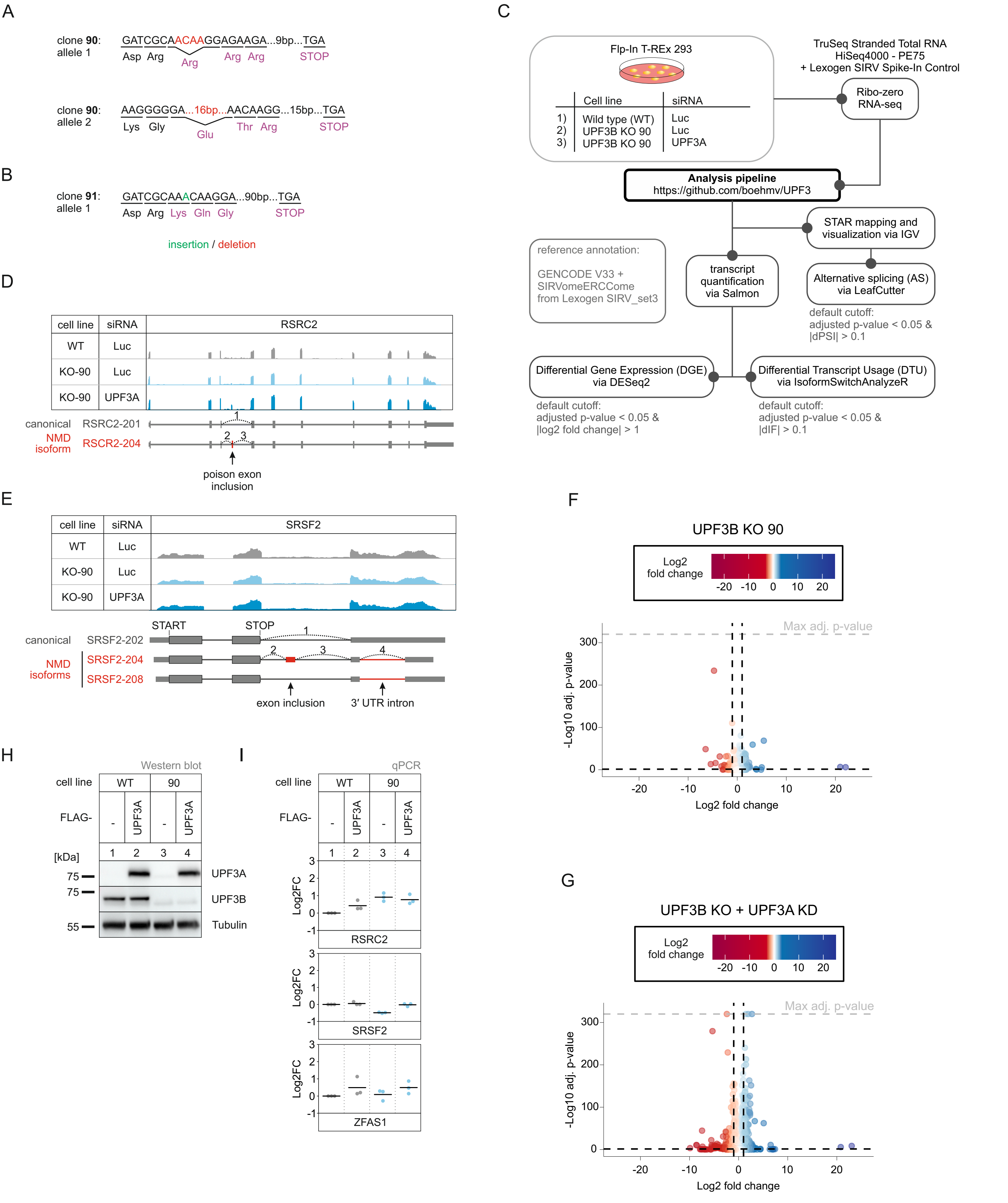


**Figure EV3 - No NMD inhibition by UPF3A in the absence of UPF3B**

**A,B** The targeted exon region and anticipated PTC location following insertions (green) or deletions (red) are indicated for detected alleles of UPF3B in KO clone 90 (A) and 91 (B).

**C** Schematic overview of the analysis pipeline.

**D,E** Read coverage of RSRC2 (D) and SRSF2 (E) from the indicated RNA-seq sample data with or without UPF3A siRNA treatment shown as Integrative Genomics Viewer (IGV) snapshot. The canonical and NMD-sensitive isoforms are schematically indicated below.

**F,G** Volcano plots showing the differential gene expression analyses from the indicated RNA-Seq datasets. The log2 fold change is plotted against the -log10 adjusted p-value (padj). P-values were calculated by DESeq2 using a two-sided Wald test and corrected for multiple testing using the Benjamini-Hochberg method.

**H** Western blot analysis of WT and UPF3B KO cells (clone 90) with or without expression of FLAG-tagged UPF3A rescue construct. UPF3A and UPF3B protein levels were detected, Tubulin serves as control.

**I** Quantitative RT-PCR of the samples from (H). For RSRC2 and SRSF2 the ratio of NMD isoform to canonical isoform was calculated. ZFAS1 expression was normalized to C1orf43 reference. Data points and means are plotted as log2 fold change (log2FC) (n=3).

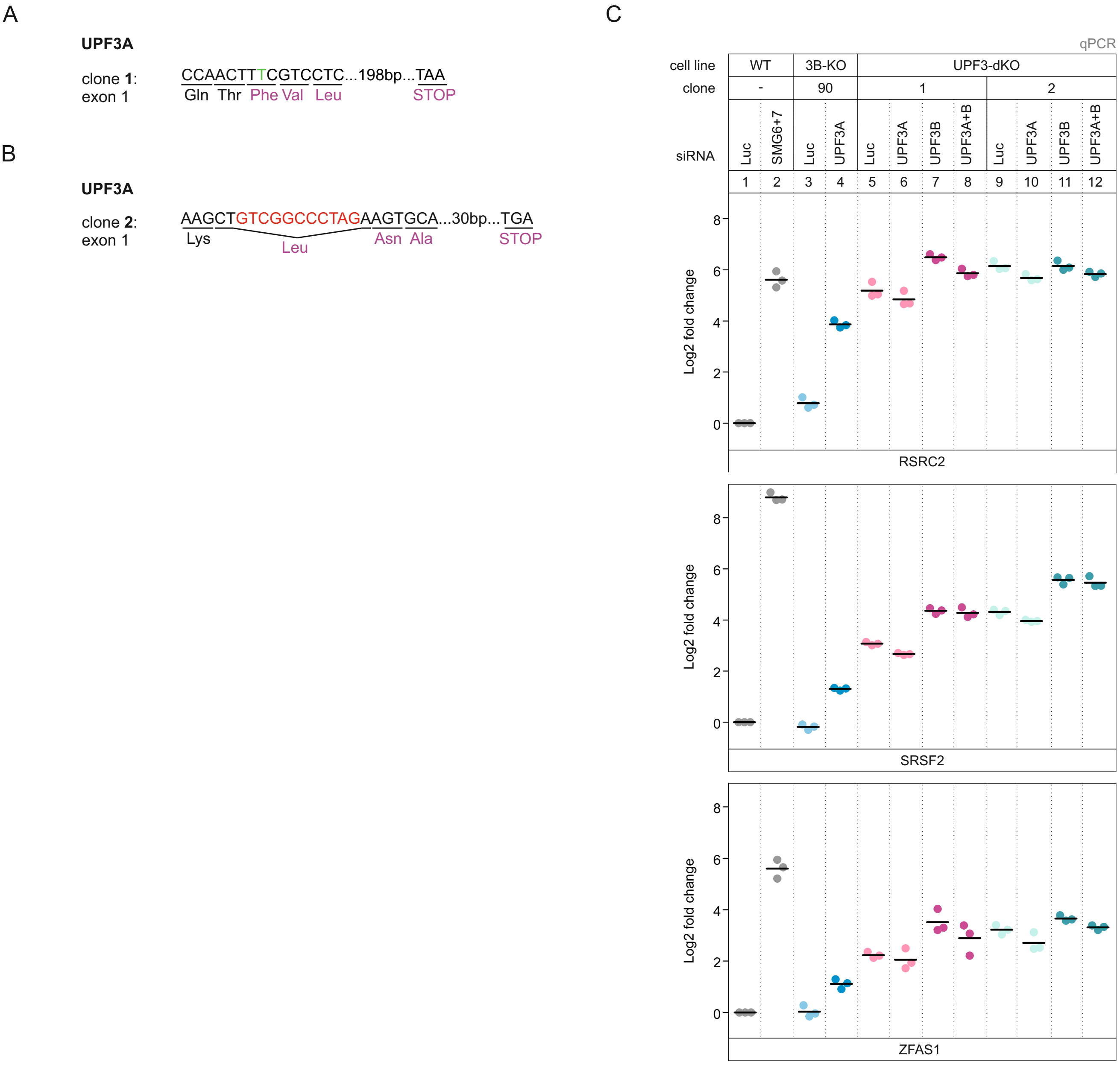


**Figure EV4 - UPF3 dKOs express residual amounts of UPF3B**

**A,B** The targeted exon region and anticipated PTC location following insertions (green) or deletions (red) are indicated for detected alleles of UPF3A in dKO clone 1 (A) and 2 (B). The UPF3B genomic locus is shown in fig EV3A.

**C** Quantitative RT-PCR of the indicated cell lines with the indicated KDs. For RSRC2 and SRSF2 the ratio of NMD isoform to canonical isoform was calculated. ZFAS1 expression was normalized to C1orf43 reference. Data points and means are plotted as Log2 fold change (n=3).

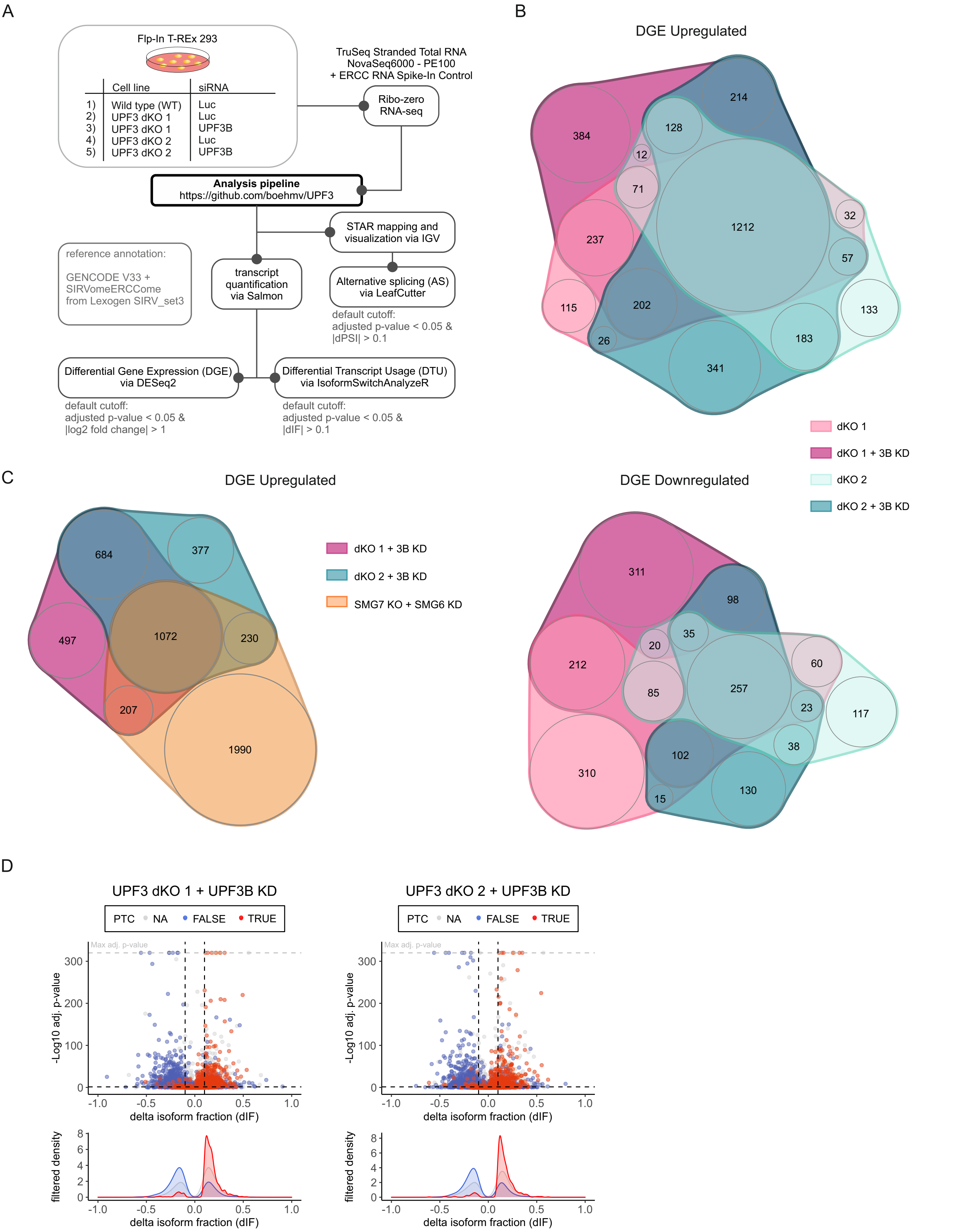


**Figure EV5 - Genes upregulated in UPF3 dKO cells are high confidence NMD targets**

**A** Schematic overview of the analysis pipeline.

**B** nVenn Diagram showing the overlap of up- or downregulated genes in the UPF3 dKO cell lines 1 and 2, both with and without a supportive UPF3B KD. Log2 fold change <-1 (downregulated) or >1 (upregulated) and adjusted p-value (padj) < 0.05. DGE = Differential Gene Expression.

**C** nVenn Diagram showing the overlap of upregulated genes in the two UPF3 dKO clones with an UPF3B KD and previously analysed SMG7 KO cells with SMG6 KD (Data ref.: Boehm et al., 2021) as control for cells with inhibited NMD. The overlap demonstrates high-confidence NMD targets. Cut-offs: log2FoldChange >1 and adjusted p-value (padj) < 0.05. DGE = Differential Gene Expression.

**D** Volcano plots showing the differential transcript usage (via IsoformSwitchAnalyzeR) in various RNA-Seq data. Isoforms containing GENCODE (release 33) annotated PTC (red, TRUE), regular stop codons (blue, FALSE) or having no annotated open reading frame (grey, NA) are indicated. The change in isoform fraction (dIF) is plotted against the -log10 adjusted p-value (adj.p-value). Density plots show the distribution of filtered isoforms in respect to the dIF, cutoffs were |dIF| > 0.1 and adj.p-value < 0.05. P-values were calculated by IsoformSwitchAnalyzeR using a DEXSeq-based test and corrected for multiple testing using the Benjamini-Hochberg method.

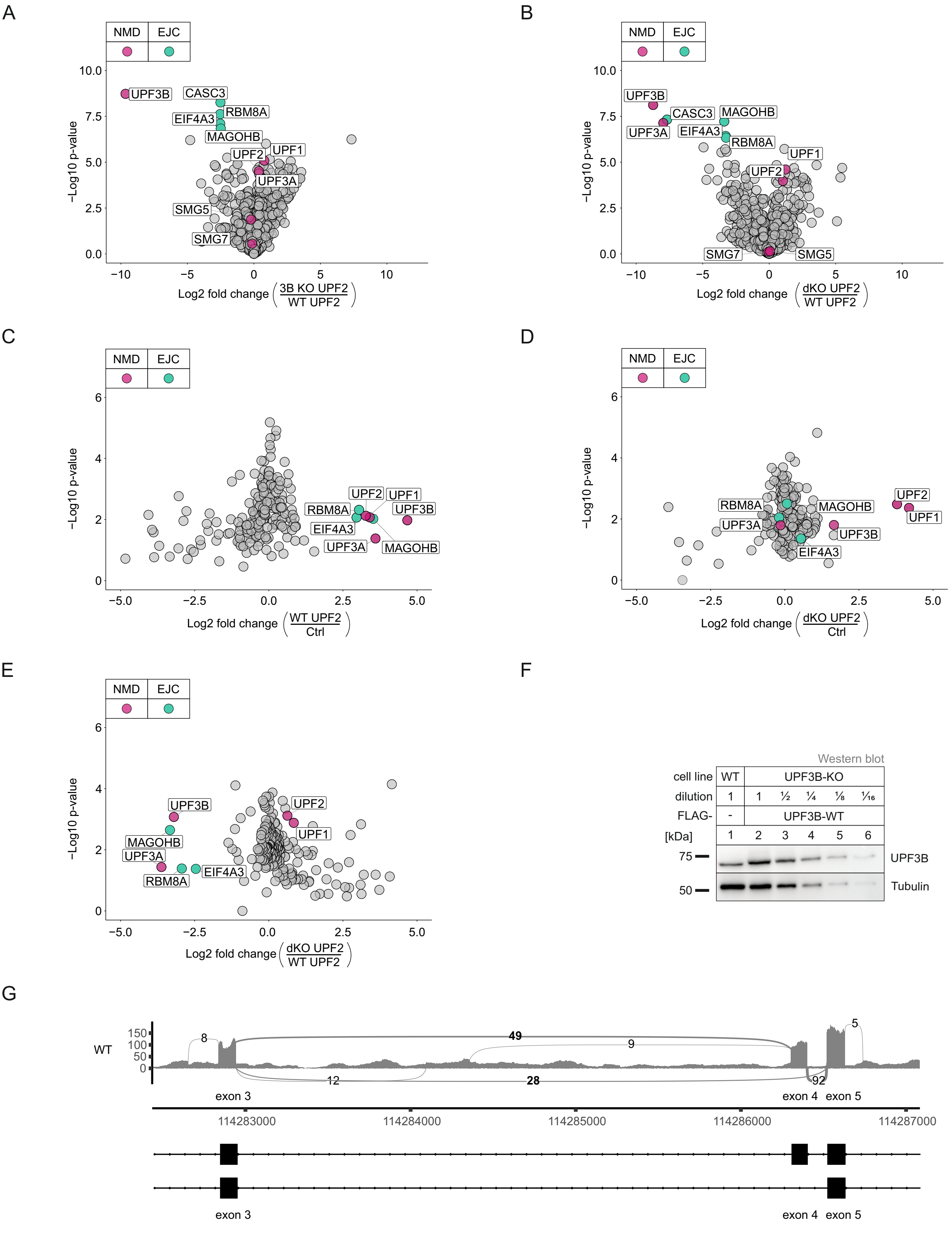


**Figure EV6 - No EJC interaction in the absence of both UPF3 paralogs**

**A,B** Volcano plots of label free mass spectrometry-based analysis of the interaction partners of UPF2 in WT cells treated with control siRNAs and the UPF3B KO clone 90 and dKO clone 1 both treated with siRNAs targeting UPF3B (n = 4 biologically independent samples). (A) FLAG-UPF2 in UPF3B KOs against FLAG-UPF2 control in WT cells, (B) FLAG-UPF2 in UPF3 dKOs against FLAG-UPF2 control in WT cells.

**C-E** Volcano plots of SILAC mass spectrometry-based analysis of the interaction partners of FLAG-UPF2 in WT cells and in the UPF3 dKO cell line 2. (C) FLAG-UPF2 against FLAG-GST control in WT cells, (D) FLAG-UPF2 against FLAG-GST control in dKO cells, (E) FLAG-UPF2 in dKO cells against UPF2 in WT cells.

**F** Western blot of WT cells and a dilution series of UPF3B KO cells expressing a UPF3B WT construct to determine exogenous expression levels. Construct expression is around 3 times higher than endogenous UPF3B.

**G** (top) Sashimi plot visualizing alternatively spliced exon and flanking exons of UPF3A in WT HEK 293 cells. Per-base expression is plotted on y-axis, genomic coordinates on x-axis. Number of junction reads are the means of three replicates (cut-off > 5). (bottom) mRNA isoforms with exons in black and introns as lines with points.
